# Superresolution microscopy reveals photoreceptor-specific subciliary location and function of Cep290

**DOI:** 10.1101/2020.10.28.357806

**Authors:** Valencia L. Potter, Abigail R. Moye, Michael A. Robichaux, Theodore G. Wensel

**Affiliations:** Verna and Marrs McLean Department of Biochemistry and Molecular Biology, Baylor College of Medicine, 1 Baylor Plaza, Houston, TX, 77030; Program in Developmental Biology, Baylor College of Medicine, 1 Baylor Plaza, Houston, TX, 77030; Medical Scientist Training Program (MSTP), Baylor College of Medicine, 1 Baylor Plaza, Houston, TX, 77030

**Keywords:** Connecting Cilium, CEP290, SIM, STORM, *Cep290^rd16^*, Y-shaped links

## Abstract

Mutations in the cilium-associated protein CEP290 cause retinal degeneration as part of multi-organ syndromic ciliopathies or as retina-specific diseases. The precise location and the functional roles of CEP290 within cilia and, specifically, the connecting cilia (CC) of photoreceptors, remain unclear. We used superresolution fluorescence microscopy and electron microscopy (TEM) to localize CEP290 in the CC and in primary cilia of cultured cells with sub-diffraction resolution, and to determine effects of CEP290 deficiency. Radially, CEP290 co-localizes with the microtubule doublets and extends beyond them. Longitudinally, it is distributed throughout the length of the CC but is strictly confined to the very base of primary cilia in hRPE-1 cells. We found Y-shaped links, the ciliary sub-structures between microtubules and membrane, at the base of the transition zone in primary cilia of epithelial cells and throughout the length of the CC. Severe CEP290 deficiencies in mouse models did not prevent assembly of cilia or cause obvious mislocalization of ciliary components in early stages of degeneration. They did not lead to loss of the Y-shaped links but caused changes in their structures. These results point to photoreceptor-specific functions of CEP290 essential for CC maturation and stability following the earliest stages of ciliogenesis.

## Introduction

Ciliopathies, genetic defects in components of primary and motile cilia, lead to a host of diseases with diverse presentations, which reflect the diverse functions of cilia (1, 2). Primary cilia, non-motile, hair-like protrusions found on virtually all mammalian cells, are elaborate structures with hundreds of protein constituents that function as cell antennae, collecting and relaying external stimuli to regulate cellular processes and maintain homeostasis. An interesting subset of ciliopathy genes can be associated either with multi-syndromic disease, often including retinal degeneration, or with retina-specific disease (3), depending on the allele, and possibly on the genetic background and other factors. Little is known about the factors that determine the precise manifestations of each ciliopathy, or about the pathophysiological mechanisms involved in retinal degeneration. The occurrence of different ciliopathy phenotypes likely depends on the precise positioning of structural and functional components.

Defects in centrosomal protein of 290 kDa (*CEP290*) are the leading genetic cause of the severe blinding condition known as Leber Congenital Amaurosis (LCA) (4–7). In addition to LCA, different *CEP290* mutations can give rise to non-syndromic retinitis pigmentosa (8) and multisyndromic disorders with associated retinal degeneration, such as Bardet-Biedl syndrome and Meckel-Gruber syndrome (5, 9–14). Understanding the functions of the protein CEP290, both in photoreceptor sensory cilia and in other primary cilia, is critical to understanding these diseases, and to guiding therapeutic interventions and genetic counseling.

Primary cilia nucleate from a mother centriole at the apical surface of the cell and have a specialized region at their base called the transition zone, beyond which extends the ciliary axoneme, a bundle of 9 + 0 microtubule doublets. The transition zone ranges from ~300 nm to 1000 nm in length and is ~300 nm in diameter, depending on cell type (1, 2, 15–18). Within the transition zone, there are filamentous structures extending from the outer surface of the axonemal microtubules to the ciliary membrane, often referred to as Y-shaped links or Y-links (the term used throughout here) because of their appearance in electron micrographs of ciliary cross-sections stained with heavy metals (19, 20). Recent electron tomographic studies suggest that their true structures in three dimensions are not very “Y-like” but confirm that they are narrower at the microtubule-end with wider bifurcations at the membrane (21, 22). The molecular composition of these structures is unknown. The ciliary membrane, while continuous with the plasma membrane, contains a unique set of membrane proteins such as ion channels and receptors (15, 23, 24). An elaborate but poorly understood network of molecular machinery within the primary cilium is necessary to ensure proper transport and distribution of ciliary components and to prevent mis-accumulation of non-ciliary proteins.

Among the numerous cell types that rely on primary cilia for proper functioning, rod and cone photoreceptor cells of the vertebrate retina carry out their entire phototransduction cascade within their cilia, making cilia essential for vision. The modified primary cilia of photoreceptors contain a light-sensing outer segment and the connecting cilium (CC), which connects the outer and inner segments. The axoneme extends from the CC into the outer segment, with microtubules extending first as doublets, and then as singlets. The light-sensing disc membranes are anchored to the axonemal microtubules. Since proteins are synthesized in the inner segment of photoreceptor cells, the CC is an essential region for the trafficking of phototransduction proteins to the outer segment. The CC is ~1100 nm in length and ~300 nm in diameter (Figure 1) (25). Inside the ring of nine microtubule doublets is the lumen, containing calcium-binding proteins called centrins (26–30), and extending from the doublets to the ciliary membrane are the Y-links (Figure 1D), which have been proposed to aid in maintaining the structural integrity of the CC (31). The distribution of Y-links along the length of the CC has not been unequivocally determined; in most ciliated cells they are restricted to the basal region of the cilia as densities that connect the axoneme and membrane, often lacking strict 9-fold symmetry. However, connections have been observed throughout the length of some primary cilia (21). The structures of the Y-links in the CC and other mammalian primary cilia appear similar, but in no cell type are their molecular components known. CEP290 has been localized to rod CC, and based on studies *in vitro* and in motile cilia (e.g., *Chlamydomonas* flagella, where it is also known as POC3), it has been proposed to play important roles in ciliogenesis, protein recruitment to the centriolar satellites, structural support of the cilium, and regulation of ciliary protein trafficking (32–34). The function of CEP290 in photoreceptors, however, remains uncertain. It has recently been proposed that retina-specific ciliopathies can arise from defects in retina-specific structures or aberrations in spatial distributions of components within the sensory cilia (outer segments plus CC) of rods and cones. In a recent study using methods similar to those employed here, we observed that a number of ciliary components, including CEP290, undergo photoreceptor-specific re-distribution in the absence of the product of another LCA-associated gene, *Spata7* (35). These results and the spectrum of CEP290-associated disease suggest there may be functions of CEP290 that are specific to retinal photoreceptor cilia.

**Figure 1.**
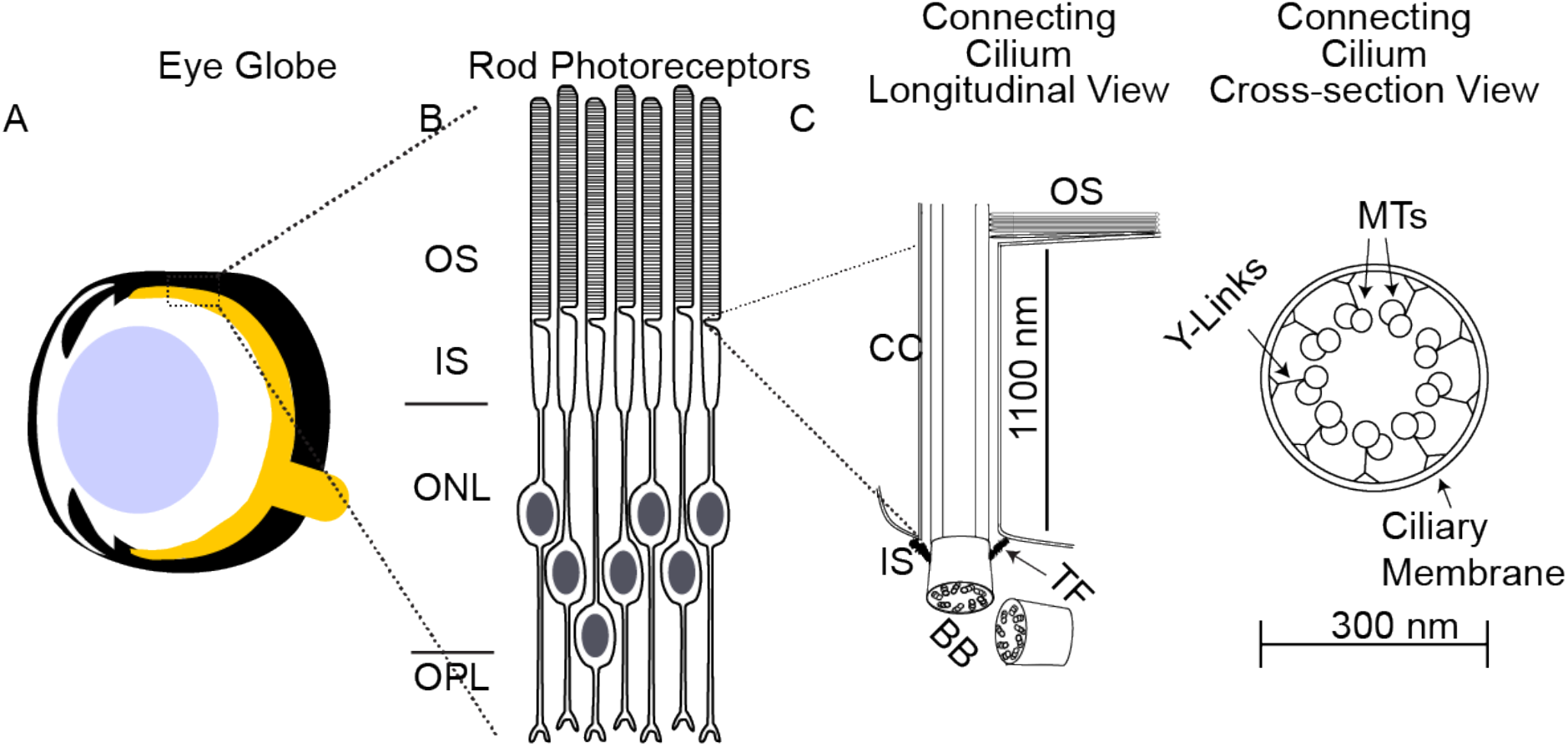
The photoreceptor connecting cilium. (A) The photoreceptors are the most posterior cells in the neural retina at the back of the eye. (B) Rod photoreceptorcells are distributed across four layers of the retina. (C) The connecting cilium links the outer segment to the inner segment. The dashed link shows the portion of the connecting cilium in the cross-sectional view. The dimensions of the connecting cilium are 1100 nm by 300 nm. OS, outer segment; IS, inner segment; ONL, outer nuclear layer; OPL, outer plexiform layer; CC, connecting cilium; BB, basal body; TF, transition fiber; MT, microtubule doublets.

A limitation of most previous studies examining localization of CEP290 in photoreceptors has been the resolution limit of conventional light microscopy. Given that the entire width of the CC is approximately 300 nm, only slightly wider than the ~250 nm full width at half maximum (FWHM) of the narrowest point-spread function practically achievable in a confocal microscope (36), imaging methods beyond the diffraction limit are needed to determine sub-ciliary distributions of CEP290 and its binding partners in wild type and *Cep290* mutant animals. Given the narrow dimensions of the cilium, we used superresolution light microscopy – structured illumination microscopy (SIM) and stochastic optical reconstruction microscopy (STORM) (37, 38) – to localize CEP290 precisely within the sub-compartments of wild type and *Cep290* mutant rod CC, as well as in primary cilia of cultured epithelial cells. We used the same techniques to examine the effects of CEP290 deficiencies on the localization of ciliary components in mouse ciliopathy models. Finally, we correlated our fluorescence localization results with new electron microscopic data revealing ciliary structures at even higher resolution.

## Results

### CEP290 localizes throughout the length of the connecting cilium and in close proximity to the microtubule doublets

To determine the spatial distribution of CEP290 within the CC, mouse retinas and retinal sections from adult animals (six weeks to eight months) were co-labeled with antibodies for CEP290 and cilia markers with known distributions. To distinguish CC staining from inner segment staining, we used CEP164, a protein whose antibodies label the transition fibers/distal appendages, structures located at the inner segment (IS)/CC interface (39). We used antibodies specific for centrins, a set of calcium-binding proteins known to be centrally localized throughout the length of the CC (26–30), to calibrate the length of the axoneme (Figure 2A). To quantify the extent of CEP290 localization along the length of the CC and to compare it to that of centrin localization, we measured the distances between the proximal and distal boundaries of CEP290 signal (Figure 2B), defined as the positions near the edges where signal was 33% of its maximal value for SIM and STORM, respectively. When measured from the IS/CC interface, marked by CEP164, the average lengths of centrin and CEP290 labeling were similar in SIM: 984 nm ± 201 nm and 1033 nm ± 200 nm, respectively. In STORM, the average lengths were 1193 nm ± 168 nm and 1336 nm ± 248 nm, respectively (Figure 2C), suggesting that CEP290’s localization may extend to the base of the outer segment, beyond the distal boundary of centrin. The small differences in length between the two imaging modalities can be explained by the differential stretching or shrinking of structures in the very different fixation and imaging media used for each method. These values are in agreement with previously reported measurements of CC length (25), and indicate that CEP290 localizes throughout the length of the CC. This finding is in contrast with its localization in other primary cilia, as discussed below, and indicates that the functional role of CEP290 in photoreceptors is performed throughout the entire CC and is not restricted to the base, as it is in the transition zones of other characterized cilia.

**Figure 2.**
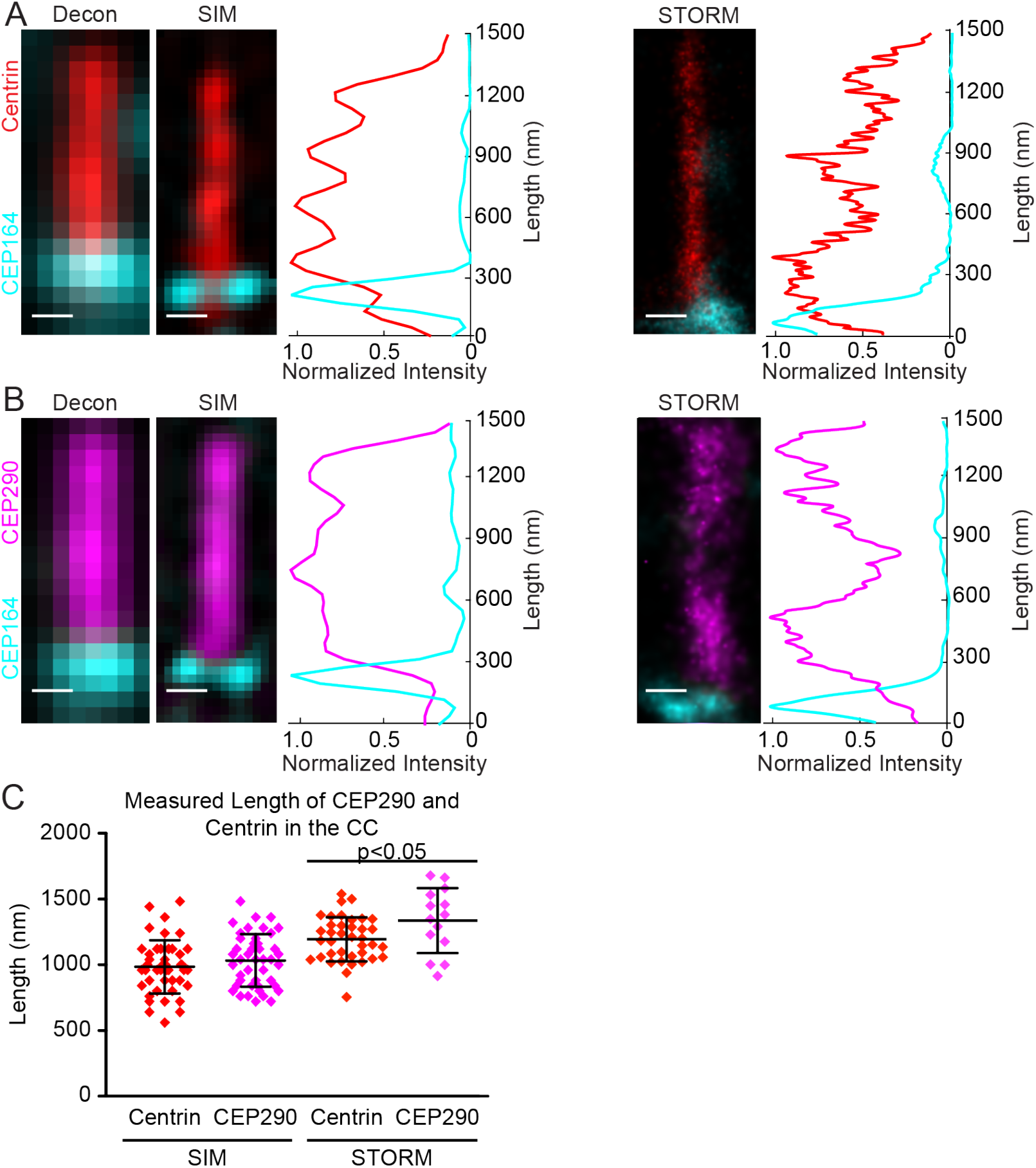
CEP29O localizes throughout the length of the CC. (A-B) Deconvolved widefield (left), SIM (middle), and STORM (right) images of a representative cilium. Row-average intensity plots are shown for SIM and STORM. (C) Dot plot graph with averages and standard deviations of the length of CEP29O and centrin in the CC for SIM and STORM. Cilia were imaged from three non-littermate mice. 33% of the maximum intensity value of each channel was used as the boudary criterion for the measurement of each cilium. Measurements were compared with Student’s t-test. CC, connecting cilium.

To determine the radial distribution of CEP290, we co-labeled retinas with antibodies for centrins, which localize to the central-most compartment of the cilium, acetylated α-tubulin (AcTub), which labels the microtubule doublets of the axoneme (40), and wheat germ agglutinin (WGA), a lectin which binds to the glycoproteins of the ciliary membrane (41). Cross sections through the CC imaged with SIM revealed that CEP290 localized outside the region of the lumen (Figure 3A). In longitudinal SIM images (Figure 3B), the width of the CEP290 signal was greater than that of centrin and AcTub signals. In STORM images, the widths of the CEP290 and AcTub signals were about the same, but wider than the centrin signal and located within the WGA-stained membrane of the CC (Figure 3C). Radial measurements of the labeled areas in longitudinal images, measured from the center of the cilium to the position of 33% of maximal signal for SIM and STORM, are shown in Figure 3D. In SIM images, CEP290 radial localization appeared significantly wider than that of centrin or AcTub. In STORM images, CEP290 radial localization was significantly wider than that of centrin; however, CEP290 localization was not significantly different from AcTub localization. The average maximal radii of centrin and AcTub were 87 nm ± 14 nm and 91 nm ± 14 nm, respectively, from SIM images, and were 66 nm ± 9 nm and 109 nm ±17 nm, respectively, from STORM reconstructions. These values are in agreement with previously reported radial measurements of the CC (25). The radius of CEP290 measured 137 nm ± 28 nm and 107 nm ± 25 nm in SIM and STORM, respectively. The small difference may reflect differences in sample preparation as suggested above, possibly leading to differential staining of CEP290 in different sub-regions of the CC in each method. Interestingly, the distribution of CEP290 signal was, in many cases, not symmetric about the central axis of the CC as would be expected for a protein in solution with a uniform concentration, but rather, was shifted either to the left or the right of the central axis. Both the width of CEP290 we observe by STORM and the asymmetry of its staining pattern are consistent with recently reported STORM images collected from cross-sectional views of centrioles in RPE-1 cells (42), but narrower than the ~250 nm width observed by STED microscopy of longitudinal views in the same cell type (18), which is more consistent with our SIM results. Figure 3E summarizes schematically the longitudinal and radial distribution of CEP290 within the rod CC. These results indicate that CEP290 localizes in close proximity to the microtubule doublets, likely in a regular structural pattern throughout the CC.

**Figure 3.**
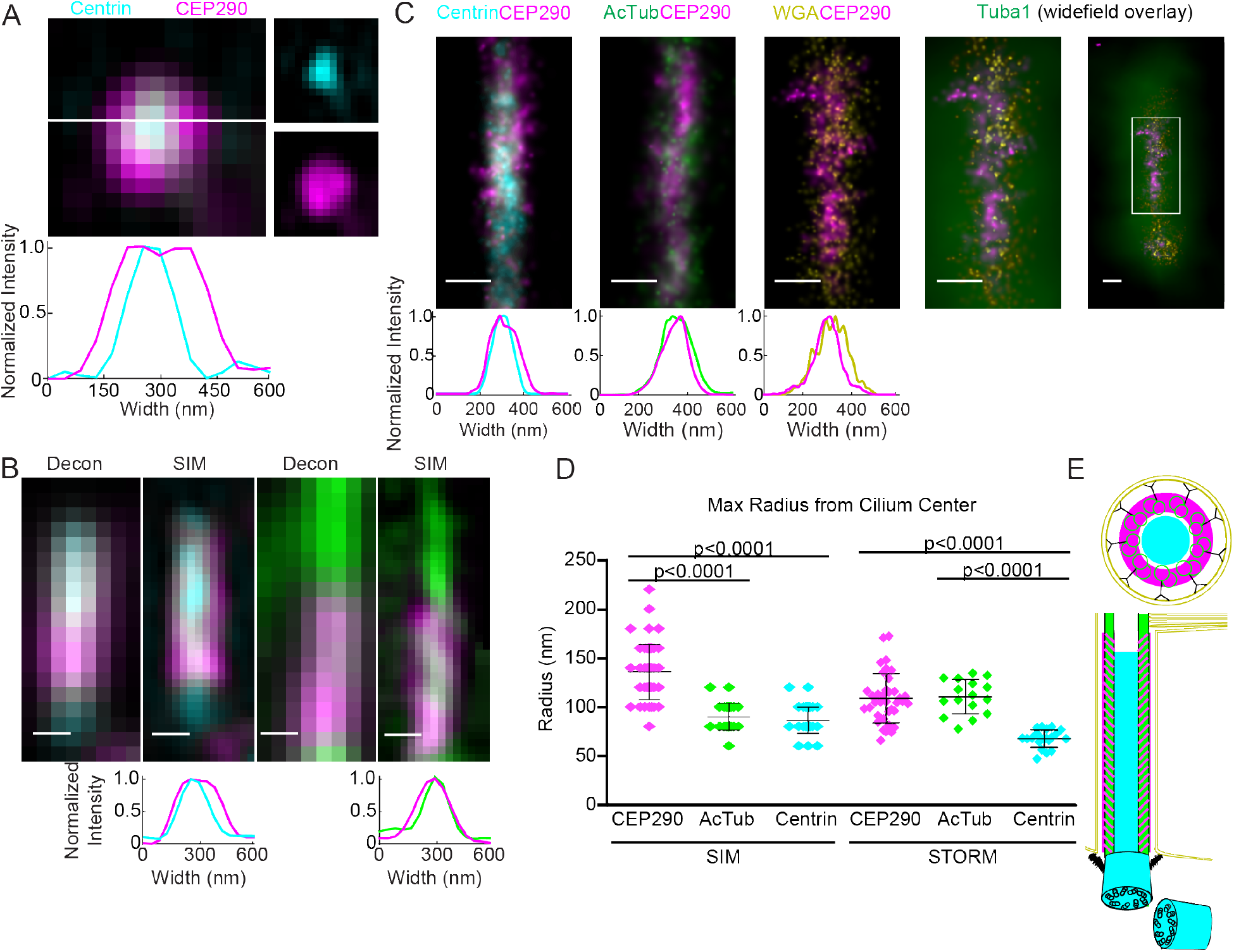
CEP29O localizes in close proximity to the microtubule doublets. (A) SIM image of a representative cross section through the CC with individual channels separated to the right. The white line depicts the position of the average intensity line plot. (B) Deconvolved widefield (left) and SIM (right) longitudinal images of a representative cilium. Row-average intensity plots are shown for SIM. Scale bar 200 nm. (C) STORM longitudinal images of a representative cilium with row-average intensity plots. For CEP29O/ WGA labelled cilia, high and low magnification widefield Tubal overlay is shown. Scale bar 200 nm. (D) Dot plot graph with averages and standard deviations of the radius of CEP29O, AcTub, and centrin in the CC for SIM and STORM. 33% of the maximum intensity value of each channel were used for the measurement of each cilium. Measurements were compared with one-way ANOVA and Tukey post-hoc test for multiple comparisons.(E) A color-coded schematic of a CC. AcTub, acetylated α-tubulin; WGA, wheat germ agglutinin; CC, connecting cilium.

### Y-links localize throughout the length of the connecting cilium

Y-links are ill-defined fibrous structures that radiate from each microtubule doublet pair to the ciliary membrane. CEP290 has been proposed to provide structural stability to the transition zone by contributing to or forming the Y-links (43). To investigate whether the Y-links of the photoreceptor CC localize to the same subcellular compartment as CEP290, *i.e*. throughout the length of the CC and in the region between the microtubules and the membrane, we performed transmission electron microscopy on sections cut as nearly perpendicular as possible to the ciliary axes. We imaged multiple cross-sectional CCs near to or overlapping with the base of the outer segment (Figure 4A). As expected, our images show Y-links in the proximal CC, identified by the absence of discs and outer segment membrane (Figure 4B-C). We also found that Y-links are present in the distal CC, a plane identified by the presence of outer segment discs *en face*, indicating that the sections were not cut obliquely. Present within the same plane were examples in which the fusion of the CC membrane with the outer segment membrane can be seen (Figure 4D). Thus, our data show that Y-links are present throughout the length of the CC, consistent with the longitudinal distribution of CEP290.

**Figure 4.**
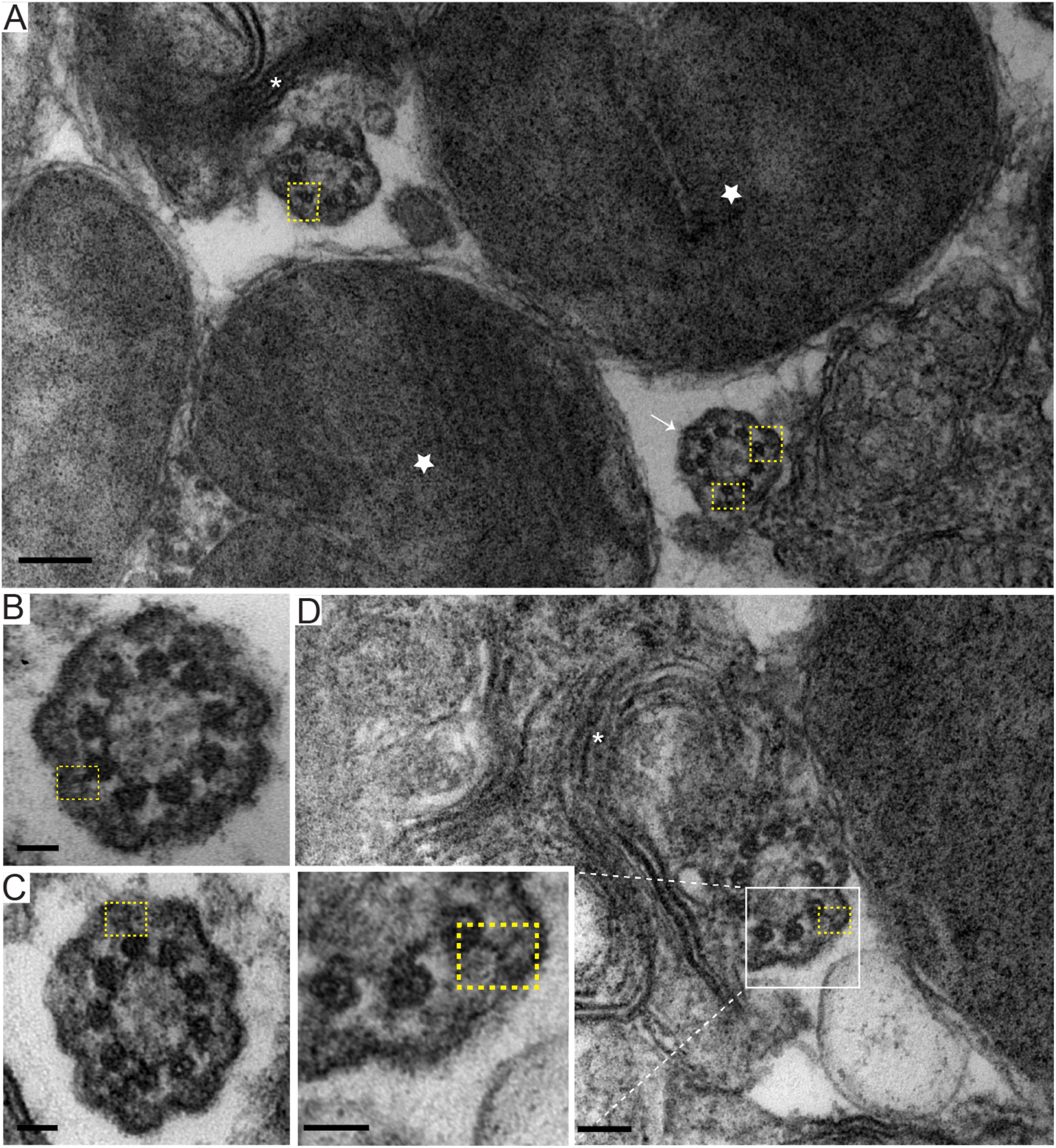
Y-links appear throughout the CC. (A) TEM image depicting the OS/CC interface. CC (white arrow), discs *en face* (star), and cilium and discs (asterisk) from a photoreceptor are shown, with MT doublets connected to Y-links outlined in yellow boxes. Scale bar 200 nm. (B-C) CC from neighboring photoreceptor cells. Y-links (yellow box) are visible. Scale bar 50 nm. (D) Image of cilium and discs (asterisk) from another photoreceptor cell. Y-links (yellow box) are visible. Scale bar 100 nm; Inset scale bar 50 nm. CC, connecting cilium; OS, outer segment.

### CEP290 localizes to the base of the ciliary transition zone in non-photoreceptor primary cilia

To compare CEP290 localization in the CC to that in primary cilia of non-photoreceptor cells, we examined CEP290 localization in primary cilia of epithelial cells. Human retinal pigment epithelium (hRPE-1) cells are cultured cells that form primary cilia upon serum starvation. Previously, it was reported that CEP290 did not localize with other transition zone proteins in hRPE-1 cells, and that, instead, CEP290 localized between the basal body and the other transition zone proteins (18). We used SIM to image ciliated hRPE-1 cells immunolabeled with centrin antibodies to identify the basal body, with acetylated a-tubulin (AcTub) antibody to label the axoneme, and with CEP164 antibodies to mark the transition fibers/distal appendages, radial structures attached to the distal end of the mother centriole in the BB (42). Consistent with previous reports (18, 32, 42), CEP290 and CEP164 localize below the base of the cilium in hRPE1 cells (Figure 5A-B). In both cases the peak of staining intensity appeared at a position more proximal than where the AcTub intensity reached its maximum value (Figure 5A-B). The average length of CEP290 labeling in primary cilia (FWHM) was 166 nm ± 78 nm (Figure 5C), and the average length for CEP164 was 198 nm ± 85 nm (Figure 5D). On average, the distance between the distal edge of CEP290 (FWHM) and the distal edge of centrin was 62 nm ± 42 nm (n=20) (Figure 5E), while the distance between the proximal edge of CEP290 (FWHM) and the proximal edge of AcTub, was 74 nm ± 147 nm (n = 33) (Figure 5F), leaving 104 nm and 92 nm, respectively, of CEP290 signal overlapping with centrin and AcTub. The average extent of CEP164 beyond centrin was 8 nm ± 59 nm (n=10); *i.e*., effectively 0 (Figure 5E). The distance between the proximal borders of CEP164 and AcTub was 140 nm ± 88 nm (n = 28) (Figure 5F), leaving 190 nm and 58 nm, respectively, of CEP164 signal overlapping with centrin and AcTub. Figure 5G illustrates schematically CEP290 localization in hRPE-1 primary cilia in relation to centrin, CEP164, and AcTub. These results suggest that CEP290 in the primary cilia of epithelial cells is predominantly located at the base of the axoneme, overlapping with but partially distal to CEP164 and centrin, and does not extend throughout the entire transition zone, consistent with previous results obtained by superresolution fluorescence(18). These results are strikingly different from those obtained in rod cells, supporting the idea of photoreceptor-specific functions for CEP290 in the CC. These results are also consistent with the hypothesis that CEP290 is associated with the Y-links, because, as shown in a recent electron tomographic study of primary cilia in a different epithelial cell line (IMCD3 cells), structures corresponding to Y-links were observed only within the first one hundred nanometers of the transition zone (21), although various structures connecting the axoneme and membrane were found beyond that point.

**Figure 5.**
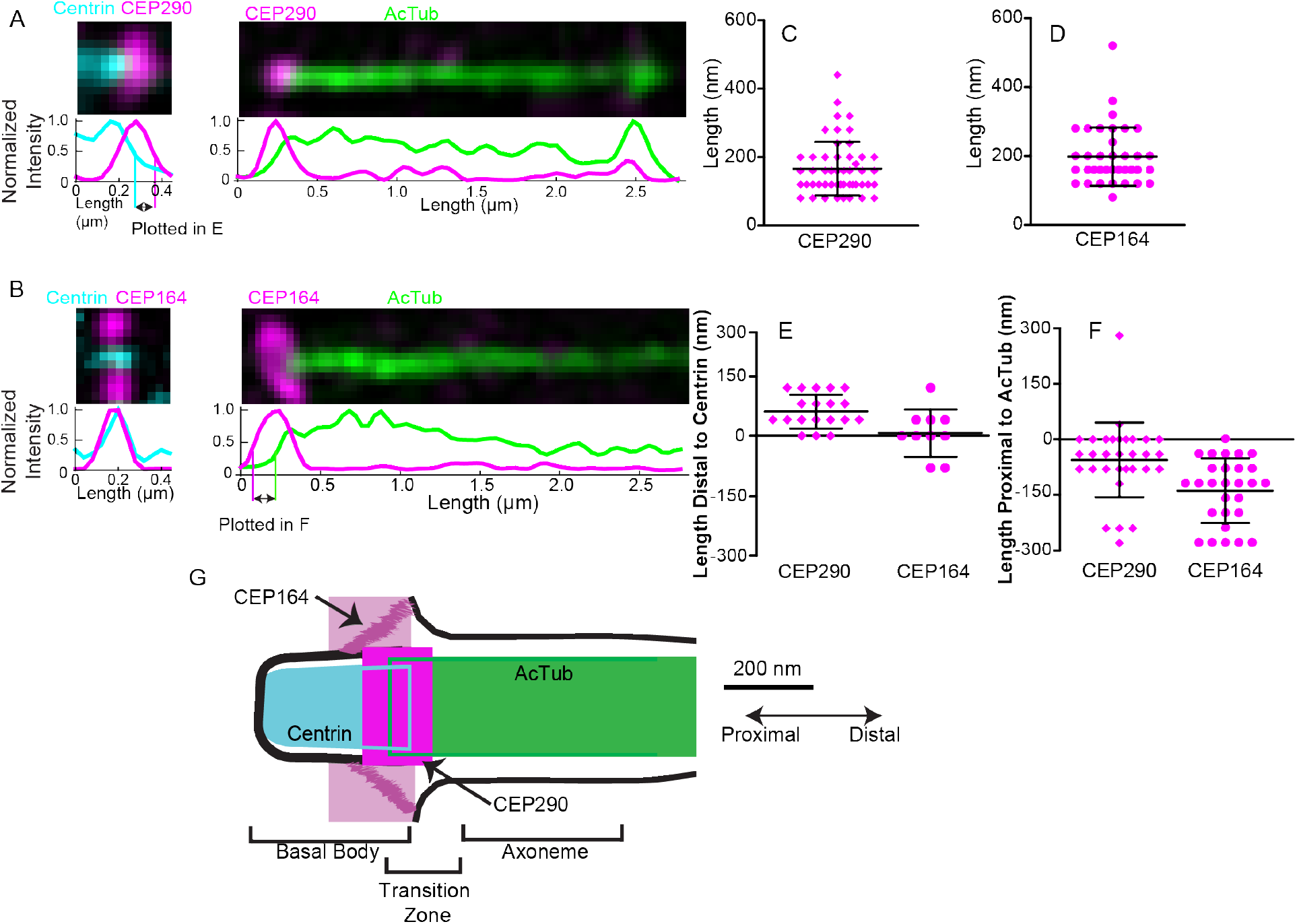
CEP290 localizes to the base of the primary cilium. (A-B) SIM images of a representative centriole and cilium for each labeling condition, CEP290 and CEP164, respectively. Row-average intensity plots are below each image. (C-D) Dot plot graph with average and standard deviation of the length of CEP290 and CEP164 in hRPE-1 primary cilia. Full width at half maximum (FWHM) intensity value of the channel was used to determine the edges for the measurement. (E-F) Dot plot graph with average and standard deviation of the length distal to centrin (right edge of centrin to the right edge of CEP290 or CEP164) or the length proximal to AcTub (left edge of AcTub to the left edge of CEP290 or CEP164). FWHM was used for the measurements. (G) A color-coded schematic of a primary cilium with results shown. AcTub, acetylated α-tubulin.

### The deletion in CEP290^rd16^ does not prevent proper CEP290 localization within the connecting cilium

We next asked how mutations in *Cep290* affect CEP290 protein localization in the CC. Leber Congenital Amaurosis (LCA) is a non-syndromic retinal disease that results in blindness or severe visual impairment in humans within the first year of life. One intronic *Cep290* mutation that leads to insertion of a cryptic stop codon and protein truncation at position 998, accounts for roughly 20% of LCA cases (6). *Cep290^rd16^* mice also display signs of a non-syndromic retinal disease and, thus, are commonly used as a model for LCA (44, 45). The *Cep290^rd16^* allele contains an in-frame 300 amino acid deletion that overlaps with the putative microtubule binding domain of CEP290 (46). *Cep290^rd16^* mice undergo rapid photoreceptor degeneration and develop abnormal, rudimentary outer segments prior to degeneration.

To determine whether the deletion affects localization of CEP290 or other CC proteins, we assessed age-matched *Cep290^rd16^* and wild type animals using superresolution microscopy. Since photoreceptor discs begin to form at ~ postnatal day 7 (P7) and *Cep290^rd16^* animals undergo photoreceptor cell death as early as P19, we used P10 animals to assess CC protein localization prior to degeneration. Surprisingly, each of the CC markers we tested displayed normal localization in the *Cep290^rd16^* rod cilia compared to wild type. Centrin localized throughout the CC lumen in wild type and mutant cilia (Figure 6A). AcTub, which labels the microtubule doublets of the CC and the outer segment in wild type rod cilia (Figure 1B-C, Figure 6B), was similarly localized in the CC and the rudimentary outer segment in the *Cep290^rd16^* cilia (Figure 6B). In both wild type and *Cep290^rd16^* P10 animals, mutant CEP290 localized longitudinally throughout the CC and radially between the axoneme and the ciliary membrane (Figure 6A-B), indicating the missing domain is not essential for CEP290c localization.

**Figure 6.**
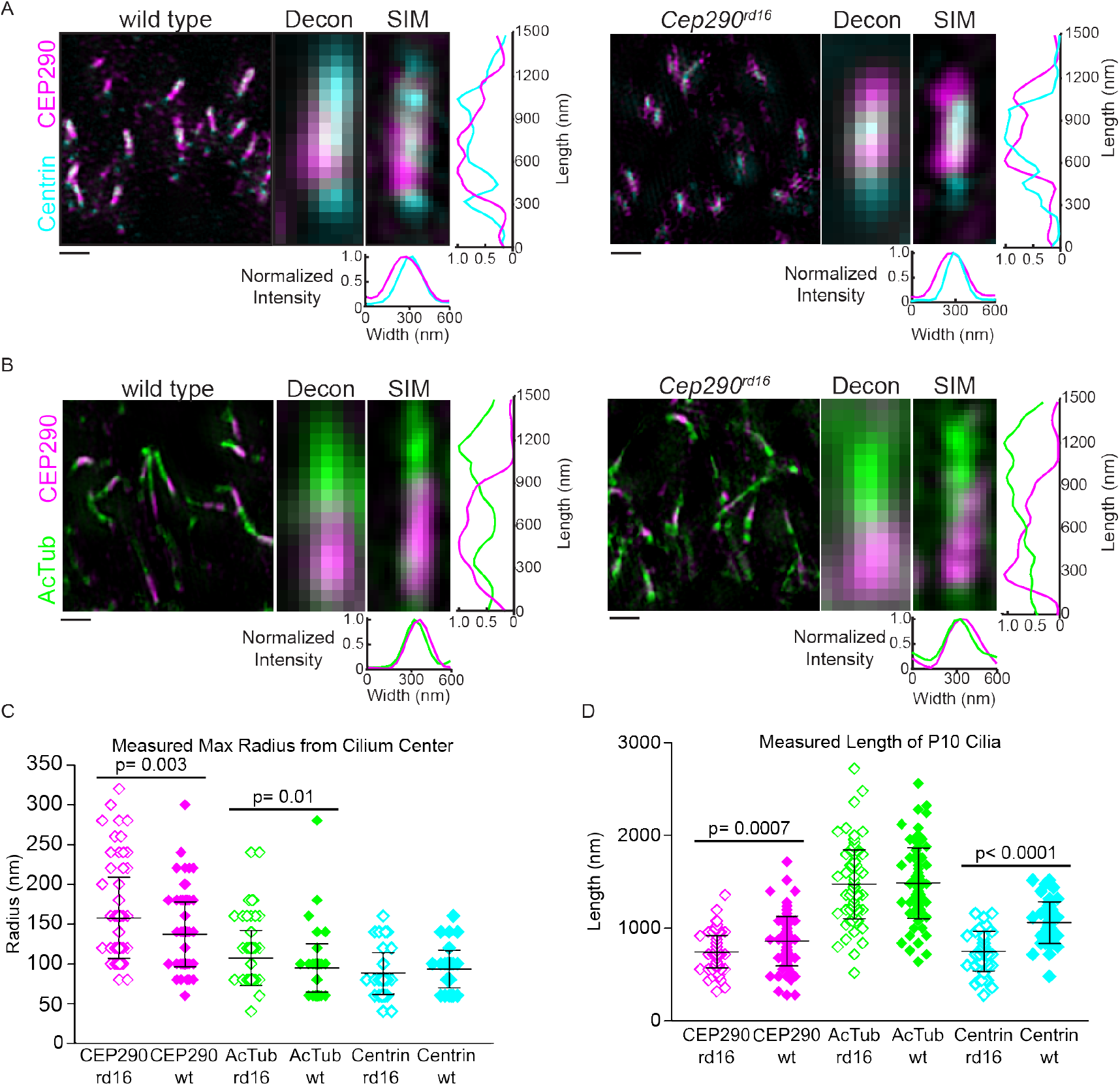
The rd16 muation in Cep290 alters CC dimensions but does not prevent proper ciliary localzation of CEP290. (A-B) SIM low magnification images (left) of cilia from wild type and rd16 animals. Scale bar 1μm. Deconvolved widefield (middle) and SIM (right) images of a representative cilium. Row-average intensity and column average intensity plots are shown for SIM. All channel intensity levels were adjusted to subsaturation for image presentation. (C-D) Dot plot graphs with averages and standard deviations of the maximum radius and length of CEP290, AcTub, and centrin in the cilium. Cilia were imaged from three non-littermate mice. 33% of the maximum intensity value of each channel was used as the boundary criterion for the measurement of each cilium. Measurements were compared with Student’s t-test. AcTub, acetylated ci-tubulin; Decon, deconvolved; CC, connecting cilium.

To compare quantitatively the distributions of centrin, AcTub, and CEP290, we measured the length and radius of antibody labeling in mutant and age-matched wild type controls. Although P10 animals have developed cilia, ciliogenesis is not yet complete, and the ciliary dimensions are not necessarily identical to those of adult mice. There was no significant difference in the average radius of centrin labeling in mutant and wild type cilia, 87 nm ± 26 nm and 93 nm ± 23 nm, respectively (Figure 6C). For both AcTub and CEP290, there are small but significant differences between the radial measurements from *Cep290^rd16^* and wild type rod cilia (Figure 6C). The AcTub radius for *Cep290^rd16^* was 107 nm ± 35 nm and for WT was 95 nm ± 30nm. The CEP290 radius for *Cep290^rd16^* was 157 nm ± 31 nm and for WT was 137 nm ± 51 nm. Length measurements revealed no statistically significant differences in the average length of AcTub in *Cep290^rd16^* and wild type cilia: 1479 nm ± 372 nm vs. 1488 nm ± 380 nm. For CEP290, there were again small but significant differences between the mutant and wild type lengths: 750 nm ± 175 nm vs. 865 nm ± 268 nm. However, there was a more substantial difference in the lengths of centrin staining between the genotypes, 763 nm ± 208 nm and 1063 nm ± 227 nm, for *Cep290^rd16^* and wild type respectively (Figure 6D) (*P* value <0.0001).

Since the rd16 deletion in CEP290 protein affects the putative microtubule binding domain of CEP290 (46), we asked whether the localization of CEP290 in relation to AcTub is affected in the *Cep290^rd16^* cilia. The average distances between these antigens in the two genotypes differed from one another by less than the pixel size of 20.025 nm (50 WT cilia and 48 *Cep290^rd16^* cilia).

### The rd 16 deletion does not prevent protein localization or substantially reduce CEP290 protein levels in Cep290^rd16^ cilia

Since CEP290 was not mislocalized in the *Cep290^rd16^* animals, we also assessed the localization of the CEP290 binding partner NPHP5. NPHP5/IQCB1 is a causal gene of LCA and Senior Löken syndrome (SLS) (47–49), and LCA and SLS patients with NPHP5 mutations phenocopy CEP290-LCA and CEP290-SLS cases (45). The C-terminal region of NPHP5 binds to the N-terminal region of CEP290 (50), forming a complex that, through unknown mechanisms, is proposed to regulate protein trafficking in primary cilia (50, 51).

We found that NPHP5 localizes throughout the CC and between the microtubule doublets and ciliary membrane in wild type photoreceptor cells at P10 (Figure 7A). NPHP5 localization does not appear to be affected by the deletion of the putative microtubule binding domain in Cep290^rd16^ animals (Figure 7A). For NPHP5, there was a small significant difference between the mutant and wild type radius (*P* value 0.04) (Figure 7B). However, there is no significant difference in the length of NPHP5 in the *Cep290^rd16^* and wild type CC (Figure 7C). Together, these localization measurements suggest that while there may be slight structural defects in CC structures of mutant retinas, the deleted region of CEP290 is not required for the localization of either CEP290 or its binding partner, NPHP5.

**Figure 7.**
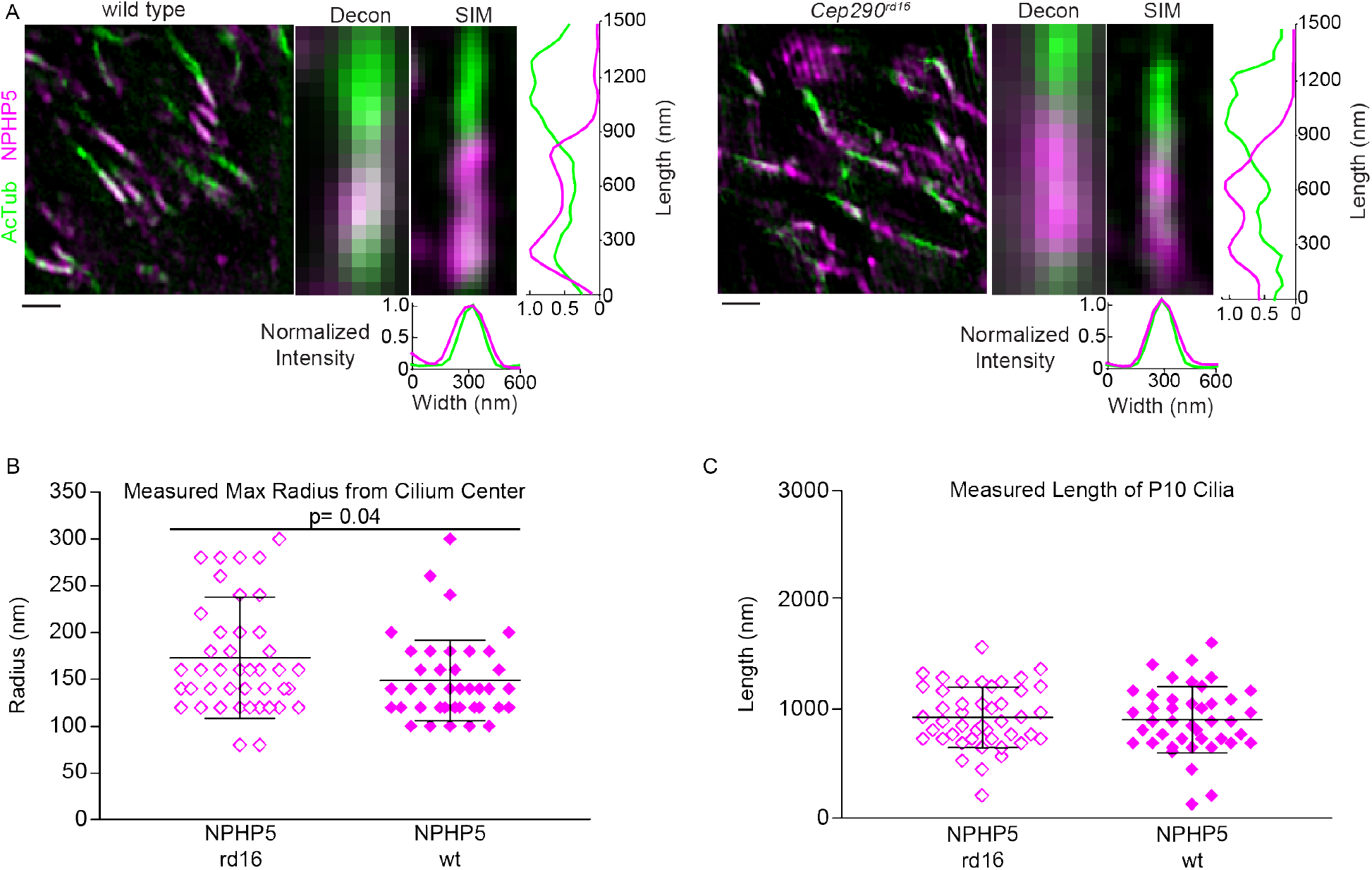
The rd16 mutation does not affect NPHP5 localization in the CC. (A) SIM low magnification images (left) ofcilia from wild type and Cep290^rd16^ animals. Scale bar 1 μm. Deconvolved widefield (middle) and SIM (right) images of a representative cilium. Row-average intensity and column average intensity plots are shown for SIM. All channel intensity levels were adjusted to subsaturation for image presentation. (B-C) Dot plot graphs with averages and standard deviations of the maximum radius and length of NPHP5 in the cilium. Cilia were imaged from three non-littermate mice. 33% of the maximum intensity value of each channel was used to determine the boundary for the measurement of each cilium. Measurements were compared with Student’s t-test. AcTub, acetylated ¤-tubulin; Decon, deconvolved; CC, connecting cilium.

In addition to NPHP5, the localization of rhodopsin, the visual pigment of the rod photoreceptor, was assessed. Rhodopsin is the most abundant protein in these cells, accounting for up to 90% of total protein in disc membranes of the outer segment (52, 53). The *Cep290^rd16^* animals form rudimentary outer segments with disordered discs (46). To determine whether outer segment localization of rhodopsin is impaired in the Cep290^rd16^ mutant prior to degeneration, retinal sections from P10 animals were costained with centrin and rhodopsin antibodies. Both wild type and *Cep290^rd16^* animals express low levels of rhodopsin at P10. While rhodopsin is able to localize to the rudimentary outer segment of *Cep290^rd16^* animals, there appears to be somewhat more rhodopsin in the inner segment and outer nuclear layer in *Cep290^rd16^* rods than in wild type rods (Supplementary Figure 1). However, the levels are too low for reliable correction of background signal and accurate quantification. These results imply that while the basic mechanisms for transporting rhodopsin to the rudimentary outer segments are maintained, there is quantitative impairment of the ability of the cells to sort rhodopsin between the inner segment and nascent discs. This deficiency, along with the shortened centrin distribution are the earliest structural or functional defects yet observed in *Cep290^rd16^* animals and may be the basis for the later defects in outer segment development and cell survival.

Since CEP290 gross protein localization appeared to be mostly unaffected by the rd16 deletion, we next asked whether the rapid retinal degeneration in Cep290^rd16^ animals could be attributed to a reduced amount of CEP290 protein. To determine whether there was a reduction in CEP290 protein product in the *Cep290^rd16^* animals, we performed immunoblots with P10 wild type and mutant retinas. Immunoblots for CEP290, AcTub, and rhodopsin demonstrate that *Cep290^rd16^* protein levels are comparable to wild type levels (Supplementary Figure 2).

### Presence of functional CEP290 is not required for connecting cilium formation or NPHP5 connecting cilium localization

Since protein localization was not grossly affected in rods from *Cep290^rd16^* pups, we investigated whether the presence of CEP290 is necessary for rod cilia formation. To address this question, we used the *Cep290^tm1.1Jgg^* animals, which serve as a model for Joubert syndrome (54). Joubert syndrome is a syndromic ciliopathy characterized by nephronophthisis, cerebellar vermis aplasia, and retinal degeneration (55). The *Cep290^tm1.1Jgg^* animals present with rapid photoreceptor degeneration and vermal hypoplasia (54). The *Cep290^tm1.1Jgg^* allele is described as a null allele; however, an alternatively spliced variant of the mutant allele may result in low residual levels of a truncated CEP290 (56). Immunostaining of retinal sections revealed little or no labeling with the CEP290 antibody in *Cep290^tm1.1Jgg^* retinas at P10 (Figure 8A-B). In both *Cep290^rd16^* and *Cep290^tm1.1Jgg^* P10 retinas, centrin and AcTub localize to the distal end of the rods, forming narrow cilium-like structures, consistent with the formation of rudimentary connecting cilia (Figure 8A-B). Furthermore, NPHP5 was correctly localized to the CC of *Cep290^tm1.1Jgg^* animals (Figure 8C) indicating that although CEP290 and NPHP5 are interacting partners within the CC, CEP290 is not necessary for the localization of NPHP5. Thus, in photoreceptors, functional CEP290 is not required for the initial formation of the CC.

**Figure 8.**
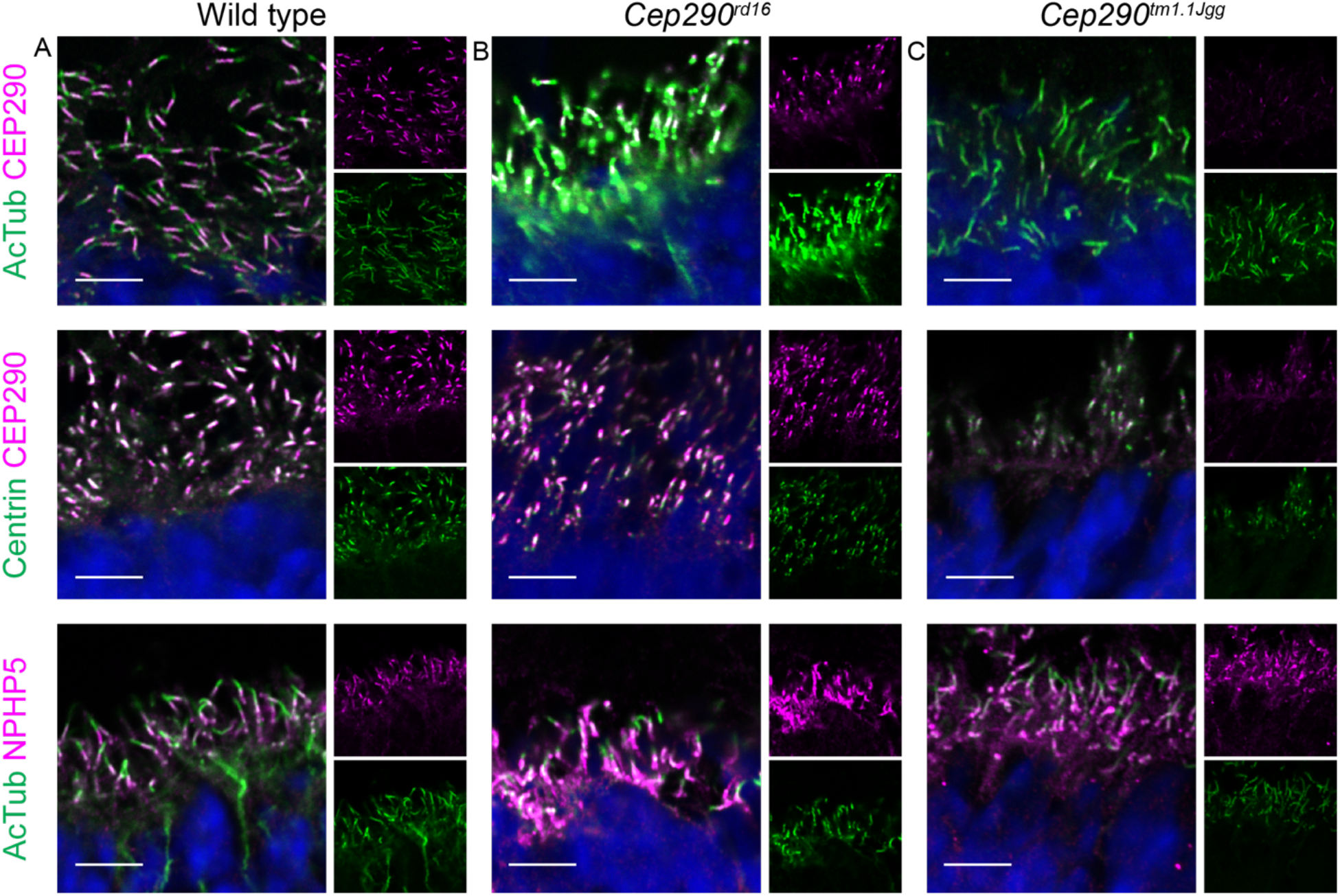
CC forms in *Cep290* mutant cilia prior to degeneration. Confocal images from P10 (A) wild type, (B) *Cep290^rd16^*, and (C) *Cep290^tm1.1Jgg^* cilia demonstrate the ability of AcTub, centrin, and NPHP5 to localize to the CC in *Cep290* mutant animals before photoreceptor degeneraton. Wild type acquisition and processing settings were applied to wild type, *Cep290^rd16^*. and *Cep290^tm1.1Jgg^* data. Scale bar 10μm. AcTub, acetylated α-tubulin; CC, connecting cilium.

### Y-links are present in Cep290 mutant connecting cilia

Since cilium-like structures labeled with the CC markers centrin and AcTub form in the presence of mutant CEP290 at either roughly normal levels (*Cep290^rd16^*) or at greatly reduced levels *Cep290^tm1.1Jgg^*, we asked whether *Cep290* mutant connecting cilia possessed Y-links by performing TEM on longitudinal and cross sections of rod photoreceptor cilia from P10 wild type, *Cep290^rd16^* and *Cep290^tm1.1Jgg^* animals. Longitudinal TEM of P10 cilia in wild type, *Cep290^rd16^* and *Cep290^tm1.1Jgg^* show that the *Cep290* mutant rods form abnormal cilia in comparison to wild type cilia (Figure 9A-C). *Cep290^rd16^* formed connecting cilia and rudimentary outer segments (Figure 9B), while *Cep290^tm1.1Jgg^* formed rudimentary connecting cilia and a small number of nascent outer segments without any obvious disc-like structures (Figure 9C). The *Cep290* mutant cilia were shorter, with reductions in the length of the outer and inner segment regions (Figure9B-C). In cross sections of the CC, wild type, *Cep290^rd16^* and *Cep290^tm1.1Jgg^* cilia have densities that radiate from the microtubule doublets and widen at the ciliary membrane in a manner similar to the Y-links (Figure 9D-F). Although the Y-links in the mutant appear grossly normal, careful measurements revealed that they are shorter (Figure 10A): 38.2 nm ± 3.8 nm in *Cep290^tm1.1Jgg^* and 40.4 nm ± 3.5 nm in *Cep290^rd16^* compared to 43.4 nm ± 3.0 nm in wild type. There were also contractions of both the axoneme and ciliary membrane (Figure 10B). The diameters for the axoneme were 171.5 ± 18.2 nm in *Cep290^tm1.1Jgg^* and 171.5 nm ± 22.9 nm in *Cep290^rd16^* compared to 187.7 nm ± 16.4 nm in wild type. The diameters for the ciliary membrane were 239.7 nm ± 23.8 nm in *Cep290^tm1.1Jgg^* and 245.8 nm ± 28.0 nm in *Cep290^rd16^* compared to 261.2 nm ± 15.7 nm in wild type. Figure 10C depicts schematically the regions measured in 10A and 10B. Note that the axoneme diameters measured by EM are the shortest widths of the cross-sections, which are different from the widths of AcTub staining measured by fluorescence of side-view images. The latter are generally wider than the actual diameter as the cilia tend to flatten somewhat in the imaging plane. Thus, the apparent widening of the AcTub staining in *Cep290^rd16^* animals may reflect a greater susceptibility to flattening, whereas the EM measurements, which indicate slight contraction of the axoneme, are likely more reflective of the relative diameters *in vivo*. There were also significant differences between the wild type and Cep290 mutant axoneme and ciliary membrane diameters. Thus, CEP290 is not required to form the Y-links and can be ruled out as the major structural protein of the Y-links. However, CEP290 is required for the Y-links to form with the correct dimensions, and therefore may be a component of the structure or required in some indirect way for their proper assembly.

**Figure 9.**
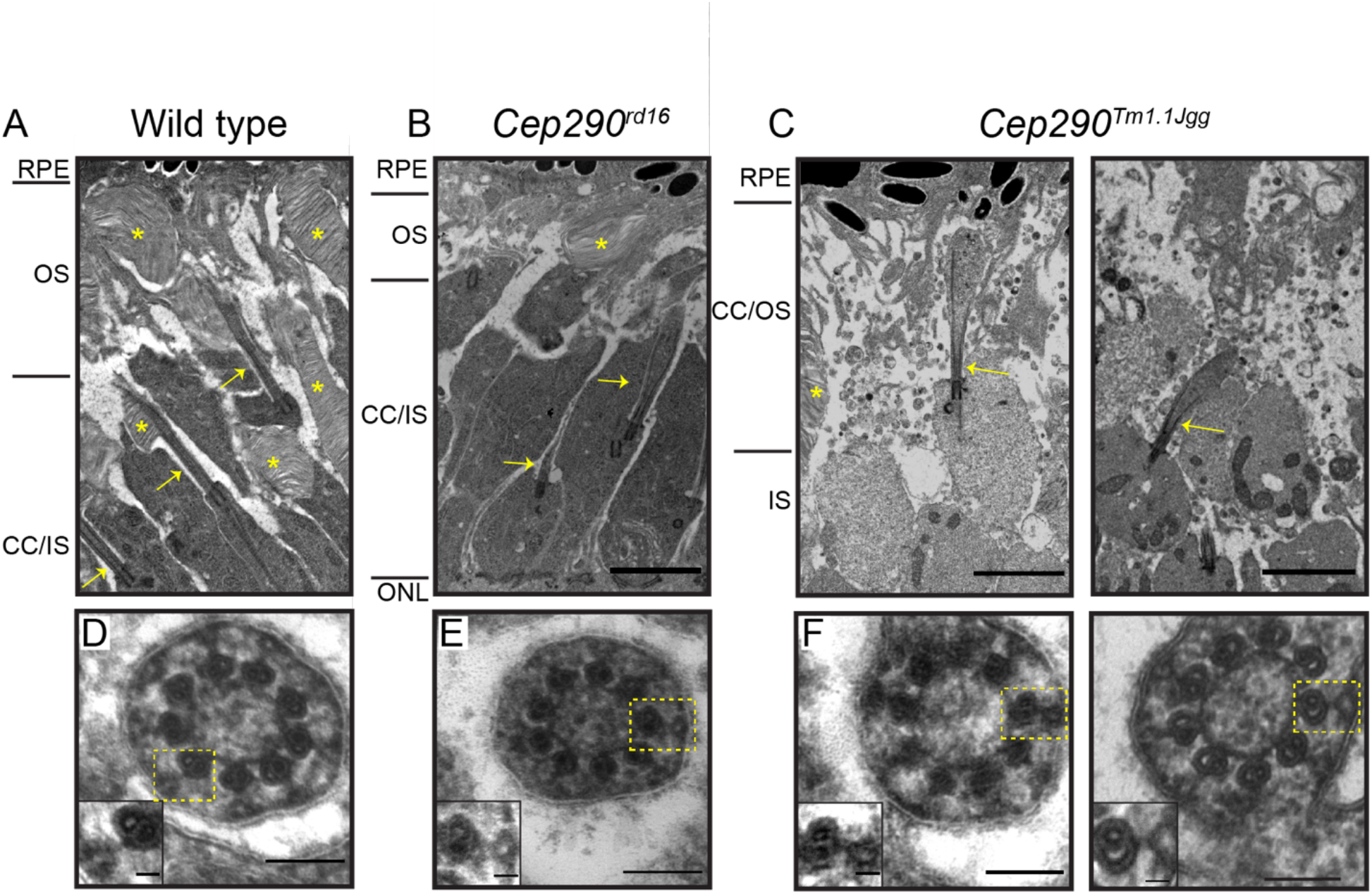
Y-links are present in P10 Cep290 mutant CC. (A-C) TEM longitudinal images of P10 photoreceptor cilia from (A) wild type, (B) *Cep290^rd1s^*, and (C) *Cep290^tm1.1Jgg^* animals depicting the (A) properly developed CC and OS, (B) rudimentary OS, and (C) rudimentary CC. CC (yellow arrow) and OS discs (yellow asterisk) are highlighted. Scale bar 2μm. (D-F) Cross section images through the CC of (D) wild type, (E) *Cep290^rd16^*, and (F) *Cep290^tm1.1Jgg^* animals. Y-links (yellow box) are visible in the wild type and *Cep290* mutants. Scale bar 100 nm. Insets highlight Y-links within the box. Scale bar 2Onm. RPE, retinal pigment epithelium; OS, outer segment, CC, connecting cilium; IS, inner segment; ONL, outer nuclear layer.

**Figure 10.**
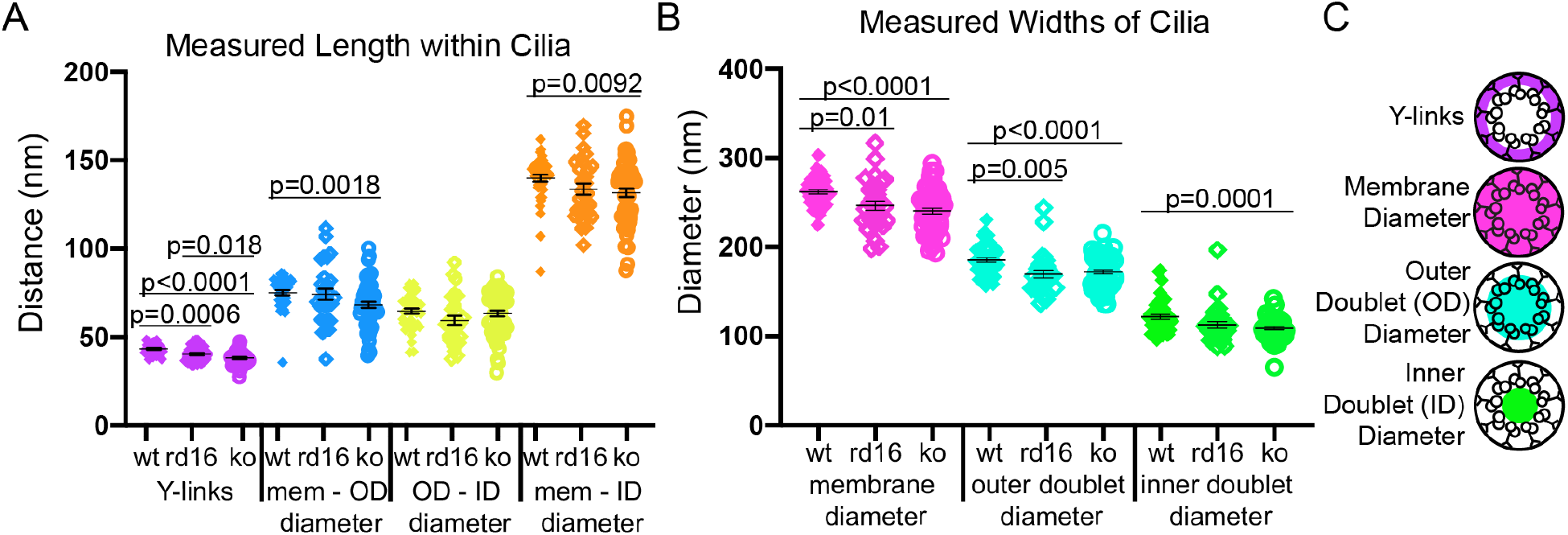
Y-link diameters are smaller in *Cep290* mutant photoreceptor CC. (A-B) Dot plot graphs with averages and SEMs of the Y-links, the differences between the compartments, membrane, and outer doublet and inner doublet diameters from wild type, Cep290^rd16^, and Cep290^tm1.1.Jgg^ cross-sections. Measurements were compared with Welch’s t-test. (C) A color-coded schematic of the regions measured in A and B.

## Discussion

The results presented here address the localization of CEP290 and its influence on the localization of other proteins in rod cilia and have important implications for its function and for the mechanisms of disease caused by CEP290 deficiencies. The observation that CEP290 is distributed throughout the length of the rod CC yet is confined to a narrow region of < 100 nm at the base of epithelial cell cilia, suggests its function may also be different in these distinct cell types. Its close association with the axoneme is consistent with proposals for its contributing to structural connections between microtubules and the membranes, as is the frequently asymmetric distribution of CEP290 with respect to the central axis of the cilium.

Based to some extent on lower resolution localization methods, there have been suggestions that what distinguishes the CC from the proximal portions of other cilia is primarily its length-that it could be considered simply a long transition zone. However, this interpretation would be an oversimplification of the multiple unique features of the CC. Whereas CEP290 is located throughout the length of the CC, it is not located throughout the transition zone of other cilia, either in primary cilia, such as the hRPE-1 cells studied here and by others (18, 42), or motile cilia, such as those in *Chlamydomonas reinhardtii* (57), but is rather confined to a narrow sub-compartment of the transition zone, and could be considered a component of the distal end of the mother centriole in those cells. Interestingly, in motile cilia of *Paramecium* (58), CEP290 was reported to reside in a distal sub-compartment of the transition zone, in close proximity to RPGRIP1L and TMEM107, but more distal than other TZ components NPHP4 and TMEM216, and to facilitate ciliary shedding induced by Ca^2+^ and ethanol.

Centrins are not observed in transition zones of hRPE-1 cell primary cilia (Figure 5), but are found throughout the length of the CC. Some essential CC-resident proteins such as RPGR^orf15^ (59) and RPGRIP1 (60) localize to centrioles in other cell types (61) and others, such as SPATA7 (35), are missing or non-essential in TZ of other cilia.

In this study, we used two different super resolution imaging techniques - SIM and STORM - to localize CEP290 and other CC proteins within the rod photoreceptor CC of adult mice. Each of these techniques, and other superresolution methods such as Stimulated Emission Depletion (STED) (18, 62) microscopy, cryo-electron tomography (25) or immunoelectron microscopy has its advantages, but all are also susceptible to distortions and artifacts, necessitating the use of multiple imaging modalities to reach reliable conclusions. While we were able to successfully image and measure protein distribution in the CC, SIM and STORM produced slightly different numbers for dimensions of labeled regions in the CC. While both techniques provide better resolution than conventional light microscopy, with STORM providing the better resolution of the two, the discrepancies between the two imaging modes can be attributed in part to different sample preparations and staining protocols; these results highlight a need for continued improvement in microscopy resolution. Advances in correlated light and electron microscopy, immunogold electron microscopy and expansion microscopy (63–66) show promise, as techniques are improved for allowing the preservation and visualization of the ultrastructure while maintaining the antigenicity necessary for immunolabeling.

Despite the major disruptions in CC structure and rapid retinal degeneration caused by the CEP290 mutations, their effects on the initial stages of cilium formation and localization of other ciliary proteins appear to be minimal, and the structural effects fairly subtle. These results are surprising, given suggestions that CEP290 is essential for ciliogenesis and proposals that it plays a major role in localizing NPHP5 and other transition zone proteins (32, 51, 67, 68). They are in stark contrast to those obtained for another LCA-associated protein, SPATA7, deficiencies in which we recently reported to cause drastic redistribution of CC proteins, including CEP290 (35). The idea that CEP290 is essential for ciliogenesis in the retina may have arisen from failure to look at early timepoints in the development of the CC as we have done in this study. Alternatively, the results we have observed may reflect presence of an alternatively spliced variant of CEP290 present in the retinas of the “near-null” mice (69).

Our results are consistent with another proposed role for CEP290, participation in the Y-link structures connecting the axoneme to the ciliary membrane (70). We find that both these Y-links and CEP290 are distributed along the length of the CC. A recent electron tomographic study of primary cilia in epithelial cells (21) demonstrated that their Y-links radiate outward from the microtubule doublets in a narrow proximal portion of the axoneme, well within the 100 nm of CEP290 labeling we observe at the base of the primary cilium in cultured epithelial cells.

However, our results are not consistent with the idea that it is a major structural component of the Y-links, as they do form in both mutants, albeit it in a quantitatively altered structure. A previous study of a CEP290 mutation in *Chlamydomonas reinhardtii* (43) reported a reduction in the number of Y-links, but not obvious abnormalities in structures of those that were present. There were also alterations in the microtubule-to-membrane distances in that mutant, but in that case the distance was increased, rather than decreased as we observe in the CC. All these results support the idea that CEP290 is a component of, or major interactor with, the Y-links and plays an important role in stabilizing the cilium but is not the major structural component bridging the axoneme and ciliary membrane. While we are now able to place additional constraints on CEP290 functions, current evidence suggests it likely has several, both those common to cilia generally, and those specific for the CC.

## Methods

### Animals

Wild-type mice used for this study were C57BL/6 aged post-natal day 10 (P10) to 8 months. *Cep290^rd16^* and *Cep290^tm1.1Jgg^* mice backcrossed to C57BL/6 were acquired from Jackson Laboratory (*CEP290^rd16^*-Stock No: 012283; *Cep290^tm1.1Jgg^*-Stock No: 013702). *Cep290* mutant animals age P10 were used for this study. For immunostaining and immunoblotting, at least 3 wild type, *Cep290^rd16^* and *Cep290^tm1.1Jgg^* mouse replicates were used. Heterozygous *Cep290^tm1.1Jgg^* crosses were bred to produce *Cep290^tm1.1Jgg^* mice. *Cep290^rd16^* animals were identified by mutagenically separated PCR using the following primers – mutant forward 5’-CCACCCCATCTTCATGTG-3’, wild type forward 5’-TGTGAAGTGAACCCATGAATAG-3’, and universal reverse 5’-CCCTCCAATATCAGGAAATGA-3’. *Cep290^tm1.1Jgg^* animals were identified by mutagenically separated PCR using the following primers – mutant forward 5’-TGGAAGACCAGGCTTCAGAG-3’, mutant reverse 5’-GGCTCACTGTGATCTTGTGC-3’, wild type forward 5’-GTAAGTGCCCGACAGCTACC-3’, and wild type reverse 5’-AGCGCAGTGCAGAGTATGTG-3’. All mouse strains were screened for the absence of *rd1* and *rd8* alleles. All procedures were approved by the Baylor College of Medicine Institutional Animal Care and Use Committee and adhered to the Association of Research in Vision and Ophthalmology (ARVO) guidelines for the humane treatment and ethical use of animals for vision research.

### Cell Culture

Human hTERT-RPE-1 (hRPE-1) cells were grown in 50/50 Dulbecco’s modified eagle’s medium (DMEM)/ F12 supplemented with 10% FBS and 10ug Hygromycin B at 37C in a humidified 5% CO2 atmosphere. To induce ciliation, cells were grown to 70% confluency, split 1:1 in normal growth media on 1.5 coverslips and allowed to settle for 12 hours prior to changing to starvation media (DMEM/F12 with 0.5% FBS). Cells remained in starvation media for 36 hours.

Cells were fixed with 2% PFA diluted in 1xPBS at room temperature for 10 minutes, blocked in PBSAT (1x PBS with 1% BSA and 0.1% Trition-X100) at room temperature for 30 minutes, incubated with primary antibody in blocking buffer for 45 minutes at room temperature, and secondary antibody in blocking buffer for 30 minutes. Coverslips were mounted with Prolong Glass (Thermo Fisher).

### Antibodies

The following commercial antibodies were used CEP290, Bethyl Laboratory, A301-659A; acetylated a-tubulin, Santa Cruz, sc-23950; NPHP5, Proteintech, 15747-1-AP; Centrin, EMD Millipore, 04-1624; CEP164, Proteintech, 22227-1-AP; CEP164, Santa Cruz, sc-515403; wheat germ agglutinin (WGA)-A647, Molecular Probes, W32466; Atto488-Tuba1, Antibodies-online.com, ABIN1169085. Rat anti-CEP290 antibody was a gift from the Swaroop laboratory. Mouse anti-1D4 was generated in house. For SIM and confocal imaging, antibodies were used at a concentration of 1 μg/150 μL. For STORM, antibodies were used at a concentration of 10 μg/mL, with the exception of rat anti-CEP290 antibody and Atto488-Tuba1 which were used at concentrations of 5 μg/mL and 7.5 μg/mL, respectively.

### Sample preparation for confocal, deconvolution, and SIM immunofluorescence microscopy

Eye cups from mice aged 4-8 months (Figures 2–5) or 10 days postnatal (Figures 6, 7, 9) were fixed for five minutes in 1% PFA (Electron Microscopy Science) diluted in 1x PBS, cryopreserved in 30% sucrose overnight at 4C, embedded in Optimal Cutting Temperature (OCT) media, and flash frozen by liquid nitrogen. Eye cups were cryosectioned at 8 μm thickness. Sections were blocked in 2% Normal Goat Serum (NGS, Fitzgerald Industries) or Normal Donkey Serum (NDS, Jackson ImmunoResearch) + 2% Fish Scale Gelatin (Sigma) + 2% Bovine Serum Albumin (Sigma) + 0.2% Triton X-100 dilute in 1x PBS and probed with 0.5 - 1 μg of primary antibody at room temperature overnight. After goat or donkey secondary antibody labeling (Thermo Fisher) (1:500), sections were mounted with Vectashield Antifade (Vector Labs) and Prolong Glass (Thermo Scientific) and covered with 1.5 coverslip (Leica) for imaging on a Leica TCS-SP5 confocal microscope. Prolong Glass mounting media was selected because the refractive index (1.520) is more compatible with SIM and reduced chromatic artifacts.

### Deconvolution microscopy and SIM imaging and image analysis

For deconvolved wide-field microscopy and SIM, sections were imaged on a DeltaVision OMX Blaze v4 (GE Healthcare) equipped with 405 nm, 488 nm, 568 nm, 647 nm lasers and a BGR filter drawer, a PLANPON6 60x / NA 1.42 (Olympus) using oil with a refractive index of 1.520, and front illuminated Edge sCMOS (PCO). For SIM, a total of 15 images were acquired per section per channel at a z-spacing of 125 nm. Deconvolved images were acquired in conventional mode, while SIM images were acquired in SI mode. Reconstructions were performed in Softworx 7 software.

Deconvolved images were deconvolved and aligned, and SIM images were reconstructed using SI reconstruction and OMX alignment. Default deconvolve and reconstruction settings were used. After analysis, reconstructions were processed in Fiji/ImageJ, and the Straighten tool was applied to straighten curved or bent cilia to acquire accurate profiles. ROI’s of digitally straightened deconvolved and STORM reconstructions were measured using row-average profiling, which plots the average intensity across the width of the ROI for each row of pixels along the length of the ROI. Pixels were converted to nm for accurate scaling. From these row-average profiles, the edges of antibody labeling in SIM were set as 33% of the maximum intensity value for connecting cilia and FWHM for hRPE-1 primary cilia, and radius was measured as the maximum intensity value for either acetylated a-tubulin (AcTub) and centrin to 33% maximum labeling for the protein of interest. All measurements were made in a 1.1 μm longitudinal region just above the basal body that corresponds to the length of the ultrastructural connecting cilium and provided as mean ± standard deviation. For radius measurements using SIM images, signals that extended beyond either the AcTub or centrin labeling were measured. All measurements were rounded to the nearest nanometer. Measurements were not collected for deconvolved images, due to its poor resolution in the focal plane.

### Transmission electron microscopy (TEM)

Adult mice were euthanized by CO_2_ asphyxiation followed by decapitation. Eye globes were enucleated and fixed in 0.1M sodium cacodylate buffer (pH 7.2) containing 2.5% glutaraldehyde at room temperature for 10 minutes. Cornea and lens were removed from the globe and the fixation of the remaining eye cup continued for 2 hours. After rinsing in buffer, the eye cups were post-fixed and heavy metal-contrasted with potassium ferrocyanide, osmium tetroxide, thiocarbohydrazide, uranyl acetate and lead aspartate. Next, the eye cups were dehydrated in acetone and embedded in Embed 812 resin. Serial block-face imaging of the resin blocks was performed on a scanning electron microscope (SEM; Mira 3, Tescan) equipped with an in chamber ultramicrotome (3View, Gatan). Serial images of the sectioned block-face (200 nm between sections) were observed on a digital monitor until the tissue plane was reached containing the outer segment-connecting cilium interface of the photoreceptors. The sectioned block was then removed from the SEM and placed in a conventional ultramicrotome for routine sectioning (80-100nm thin) and collection on copper grids (200 mesh) for imaging on the transmission electron microscope (Tecnai 12, FEI). Optimal cross-sectional images of the cilia were achieved using a goniometer specimen chamber capable of ± 60-degree tilting in conjunction with a motorized rotating (360 degree) specimen holder.

For P10 studies, mice were euthanized by CO2 asphyxiation followed by decapitation at P10. Eyes were enucleated, the cornea and lens were removed, and eye globes were placed in fixative (2% paraformaldehyde, 2% glutaraldehyde, 3.4mM CaCl_2_ in 0.2M HEPES, pH 7.4) for 24 hours, rocking at room temperature. The eyecups were then washed in 1X PBS for 5 minutes, placed in 4% agarose, and left to polymerize for 30 minutes at 4C. 150 μm sections were cut on a vibratome and subsequently stained with 1% Tannic Acid with 0.5% Saponin in MilliQ water for 1 hour, rocking at room temperature. After rinsing in MilliQ water, the sections were then stained with 1% uranyl acetate in 0.2 M Maleate Buffer, pH 6.0 for 1 hour, rocking at room temperature. The sections were rinsed in MilliQ water and dehydrated in a series of ethanol washes (50%, 70%, 90%, 100%x2) for 15 minutes each, followed by infiltration Ultra Bed Epoxy Resin. The sections were embedded in resin between two ACLAR sheets sandwiched between glass slides in a 60C oven for 48 hours. At 24 hours, the top slide and ACLAR sheet was removed and resin blocks in BEEM capsules were stamped onto each section to allow for polymerization the following 24 hours. Ultrathin silver sections were placed on copper grids and post-stained in 1.2% uranyl acetate in MilliQ water for 6 minutes, followed by staining in Sato’s Lead for 6 minutes. Sections were imaged on a Hitachi 700 electron microscope. Measurements from radial cross sections of photoreceptor CC were performed on Image J. Since the cross sections were not perfectly circular, two intersecting measurements (along the shortest and longest axes) were taken and the shortest measurement was used in the analysis for Figure 9.

### STORM immunohistochemistry and resin embedding

Retinas from 6-8 week old wild type mice were immunolabeled for STORM using a protocol we developed previously (Robichaux et al 2019). Whole retinas were stained in a solution following a two-step protocol. First, retinas were dissected unfixed in ice cold Ames’ media (Sigma) and immediately blocked in 10% NGS (Fitzgerald Industries) + 0.2% saponin (Sigma) + 1x Protease Inhibitor Cocktail (GenDepot) diluted in 1x Ames’ media for 2 hours at 4°C. Primary antibodies (5-10 μg each) were added to the blocking buffer and incubated at 4°C for 20-22 hours. Retinas were washed 3 times for 5 minutes in 2% NGS in Ames’ media on ice before secondary antibodies were added to the same buffer and incubated at 4°C for 2 hours. Secondary antibodies used (8 μg each): F(ab’)2-goat anti-mouse IgG Alexa 647 & F(ab’)2-goat anti-rabbit IgG Alexa 555 (Thermo Fisher). In WGA-Alexa647 labeling experiments, F(ab’)2-goat anti-rabbit IgG Alexa 555 (Thermo Fisher) was used for dual labeling. Retinas were washed in 2% NGS/Ames 6 times for 5 minutes each on ice and fixed in 4% paraformaldehyde diluted in 1xPBS for 15 minutes at room temperature.

Next, retinas were re-blocked in 10% normal goat serum + 0.2% saponin diluted in 1xPBS for 2 hours at room temperature. Primary antibodies (5-10 μg each) were readded to the blocking buffer and incubated for 2 days at 4°C. After this incubation, retinas were washed 4x for 10 minutes each in 2% NGS/1x PBS. The same secondary antibodies were added to the wash buffer as before (8 μg each) for overnight incubation at 4°C. Retinas were washed 6x in 2% NGS/1x PBS for 5 minutes each before postfixation in 3% formaldehyde diluted in 1x PBS for 1 hour at room temperature.

Post-fixed retinas were dehydrated in a series of ethanol washes (15 minutes each: 50%, 70%, 90%, 100%, 100%) followed by embedding steps of increasing concentrations with Ultra Bed Epoxy Resin (Electron Microscopy Sciences) to ethanol (2 hours each: 25%:75%, 50%:50%, 75%:25%, 100% resin twice). Embedded retinas were cured on the top shelf of a 65’C baking oven for 20 hours. 500 nm - 1 μm sections were cut on a UCT or UC6 Leica Ultramicrotome and dried directly onto glass-bottom dishes (MatTek 35 mm dish, No. 1.5 coverslip).

### STORM image acquisition

Immediately prior to imaging, 10% sodium hydroxide (w/v) was mixed with pure 200-proof ethanol for 45 minutes to prepare a mild sodium ethoxide solution. Glass-bottom dishes with ultra-thin retina sections were immersed for 30-45 minutes for chemical etching of the resin. Etched sections were then washed and dried on a 50°C heat block. The following STORM imaging buffer was prepared: 45 mM Tris (pH 8.0), 9 mM NaCl, oxygen scavenging system: 0.7 mg•ml–1 glucose oxidase (Amresco) + 42.5 μg ml-1 catalase (Sigma), 10% (w/v) glucose + 100 mM MEA (i.e. L-cysteamine, Chem-Impex) + 10% VECTASHIELD (Vector Laboratories). Imaging buffer was added onto the dried, etched sections and sealed with a second number 1.5 coverslip for imaging.

Imaging was performed on the Nikon N-STORM system, which features a CFI Apo TIRF 100x oil objective (NA1.49) on an inverted Nikon Ti Eclipse microscope. STORM image acquisition was controlled by NIS-Elements Ar software.

To begin a STORM acquisition, both the 561 nm and 647 nm laser lines were increased to maximum power to photobleach the fluorescence and initiate photoswitching.

Imaging frames were collected at ~56 frames per second. 50,000 frames were collected for each imaging experiment.

### STORM image analysis

2D-STORM Analysis of STORM acquisition frames was performed using NIS Elements Ar Analysis software. Analysis identification settings were used for detection of the individual point spread function (PSF) of photoswitching events in frames from both channels to be accepted and reconstructed as 2D Gaussian data points. These settings were as follows: Minimum PSF height: 400, Maximum PSF height: 65,636, Minimum PSF Width: 200 nm, Maximum PSF Width: 700 nm, Initial Fit Width: 350 nm, Maximum Axial Ratio: 2.5, Maximum Displacement: 1 pixel.

After analysis, reconstructions were processed in Fiji/ImageJ using the similar method described to analyze SIM reconstructions. The edges of STORM clusters were set as 1/e the maximum intensity value instead of ½ the maximum intensity.

### Immunoblotting

P10 retinas were collected in 2x protease inhibitor (Roche) diluted in 1x PBS (6.7mM PO_4_ with Ca^++^ and Mg^++^) (Hyclone) from wild type and *Cep290^rd16^* animals, sonicated in 1x SDS sample buffer (250mM Tris pH 6.8, 10% SDS, 30% glycerol, 5% β-mercaptoethanol), and 0.25 −1 retina per lane were loaded on 8 or 14% gels for SDS-PAGE. Gels ran in 1x Tris Glycine (TG)-SDS buffer (25mM Tris, 192mM glycine, 0.1% SDS, pH 8.3) (Bio-Rad). Proteins were transferred to unsupported nitrocellulose membranes in 1x TG (25mM Tris, 192mM glycine, pH8.3) (Bio-Rad) for 1.5 hours at 4C. Membrane was blocked in 5% skim milk for 1 hour at room temperature. Primary antibodies were added and incubated overnight in 5% non-fat dry milk at 4C. Secondary IR antibodies (LI-COR Biosciences) were incubated with the membranes for 1 hour with shaking at room temperature. Membranes were imaged on the LI-COR Odyssey. Image processing was performed in Abode Photoshop. All image contrast adjustments were applied identically to all images being compared.

### Statistics

Numbers of animals and/or replicates are indicated in each figure legend. Two-way comparisons between antigens in one genotype were analyzed by Student’s *t*-test. Multiple comparisons were analyzed by one-way Anova with Tukey’s post-hoc correction for multiple comparisons. For Fig. 10, Welch’s *t*-test was used to allow for different underlying population variances in mutants as compared to WT.

### Study Approval

All animal studies were approved by the Baylor College of Medicine Institutional Animal Care and Use Committee. They adhered to the Association of Research in Vision and Ophthalmology (ARVO) guidelines for the humane treatment and ethical use of animals for vision research.

## Author Contributions

VLP designed and executed experiments, analyzed data and wrote the manuscript. ARM and MAR designed and executed experiments, analyzed data, wrote sections of the manuscript and helped edit it. TGW supervised the project, analyzed data and helped edit the manuscript.

## Acknowledgements

We thank Melina A Agosto for technical assistance and comments on the manuscript; Michael Mancini, Fabio Stossi, Christopher Hampton, and Hannah Johnson of Baylor College of Medicine Integrated Microscopy Core (IMC) for technical support and advice, and Alan Burns, Samuel Hanlon, and Margaret Gondo of the University of Houston School of Optometry for serial-sectioning SEM and TEM.

---

**Supplementary Figure 1.**
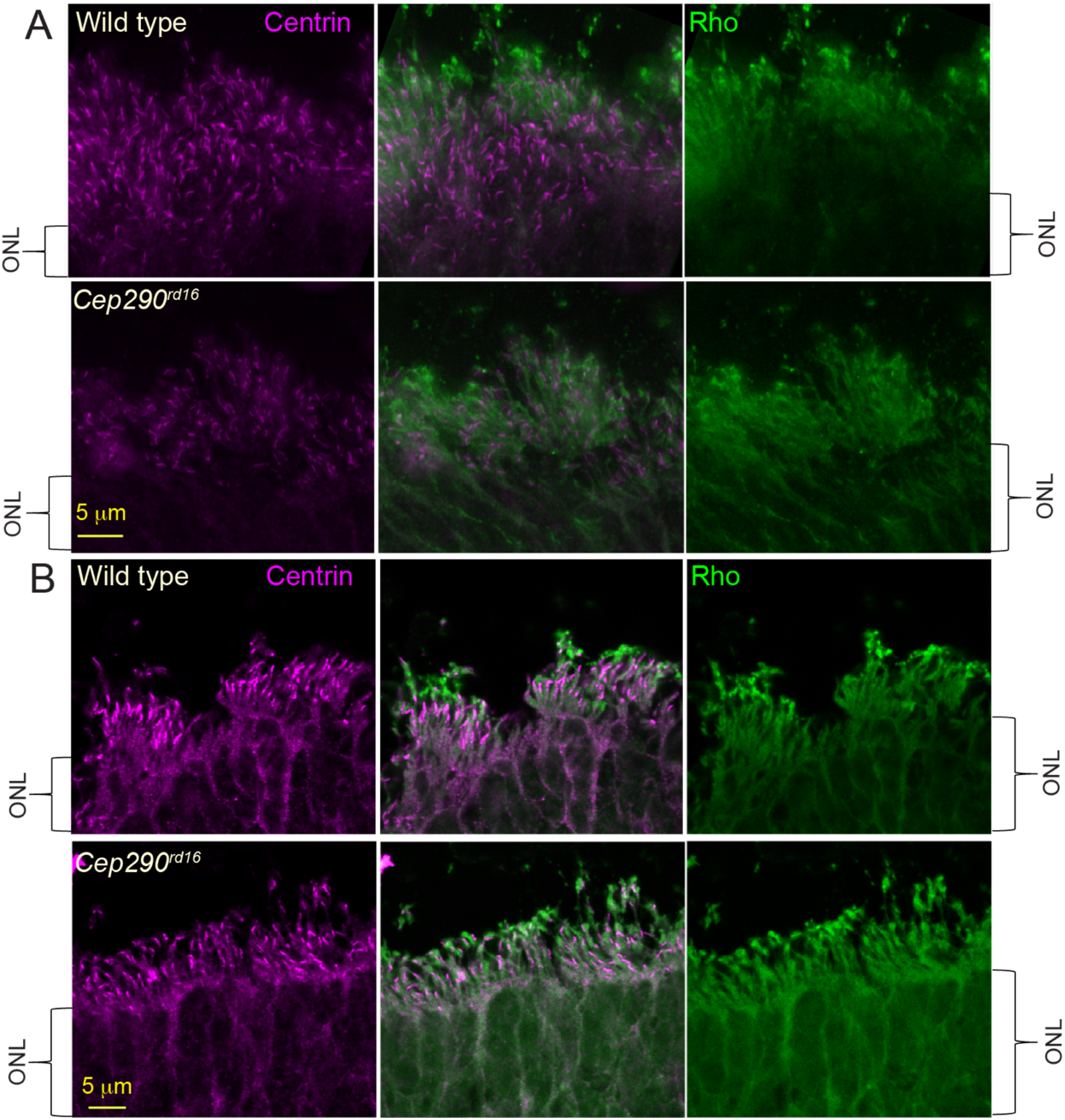
Rhodopsin localization in P10 Cep290^rd16^ cilia. (A) Deconvolved widefield and (B) confocal images of cilia from P10 wild type and *Cep290^rd16^* stained for rhodopsin (green) or centrin (magenta), marking nascent CC. Acquisition settings and image processing were identical for wild type and rd16 samples. ONL, outer nuclear layer; CC, connecting cilium.

**Supplementary Figure 2.**
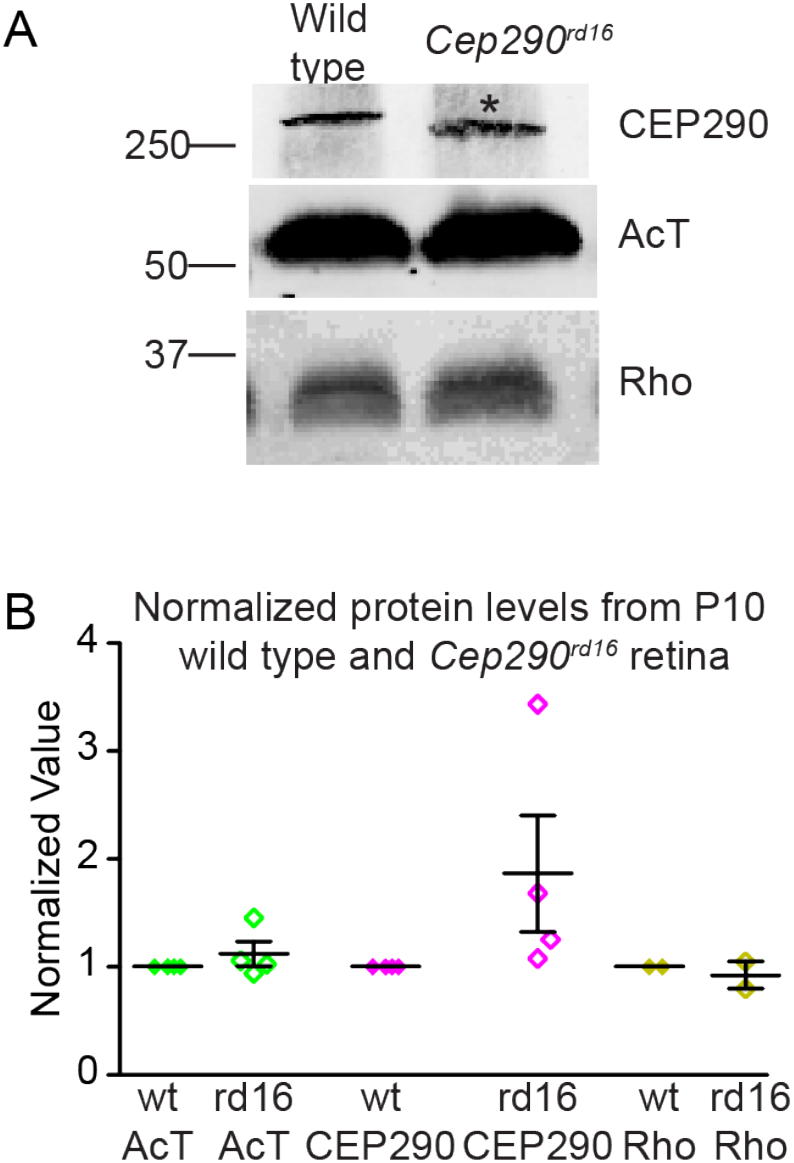
P10 wild type and Cep290^rd16^retinas have comparable protein levels. (A) Immunoblots comparing protein levels of CEP29O, AcTub, and rhodopsin in wild type and *Cep290^rd16^* retina. Expected apparent molecular weights for CEP29O, AcTub, and rhodopsin are 290 kDa, 55kDa, and 36kDa, respectively. * denotes a band shift in the *Cep290^rd16^* protein product. (B) Graph depicting relative protein levels in wild type and *Cep290^rd16^.* CEP29O blots were normalized to AcTub blots prior to normalizing CEP29O to wild type. Error bars represent SEM. AcTub, acetylated α-tubulin; Rho, rhodopsin.

## References

1. Badano JL, Mitsuma N, Beales PL, and Katsanis N. The ciliopathies: an emerging class of human genetic disorders. Annu Rev Genomics Hum Genet. 2006;7:125–48.

2. Zariwala MA, Knowles MR, and Omran H. Genetic defects in ciliary structure and function. Annu Rev Physiol. 2007;69:423–50.

3. Bujakowska KM, Liu Q, and Pierce EA. Photoreceptor Cilia and Retinal Ciliopathies. Cold Spring Harb Perspect Biol. 2017.

4. Weleber RG, Francis PJ, Trzupek KM, and Beattie C. In: Adam MP, Ardinger HH, Pagon RA, Wallace SE, Bean LJH, Stephens K, et al. eds. GeneReviews((R)). Seattle (WA); 1993.

5. den Hollander AI, Koenekoop RK, Yzer S, Lopez I, Arends ML, Voesenek KE, et al. Mutations in the CEP290 (NPHP6) gene are a frequent cause of Leber congenital amaurosis. Am J Hum Genet. 2006;79(3):556–61.

6. den Hollander AI, Roepman R, Koenekoop RK, and Cremers FP. Leber congenital amaurosis: genes, proteins and disease mechanisms. Prog Retin Eye Res. 2008;27(4):391–419.

7. Perrault I, Delphin N, Hanein S, Gerber S, Dufier JL, Roche O, et al. Spectrum of NPHP6/CEP290 mutations in Leber congenital amaurosis and delineation of the associated phenotype. Hum Mutat. 2007;28(4):416.

8. Birtel J, Gliem M, Mangold E, Muller PL, Holz FG, Neuhaus C, et al. Nextgeneration sequencing identifies unexpected genotype-phenotype correlations in patients with retinitis pigmentosa. PLoS One. 2018;13(12):e0207958.

9. Valente EM, Silhavy JL, Brancati F, Barrano G, Krishnaswami SR, Castori M, et al. Mutations in CEP290, which encodes a centrosomal protein, cause pleiotropic forms of Joubert syndrome. Nat Genet. 2006;38(6):623–5.

10. Baala L, Audollent S, Martinovic J, Ozilou C, Babron MC, Sivanandamoorthy S, et al. Pleiotropic effects of CEP290 (NPHP6) mutations extend to Meckel syndrome. Am J Hum Genet. 2007;81(1):170–9.

11. Brancati F, Barrano G, Silhavy JL, Marsh SE, Travaglini L, Bielas SL, et al. CEP290 mutations are frequently identified in the oculo-renal form of Joubert syndrome-related disorders. Am J Hum Genet. 2007;81(1):104–13.

12. Frank V, den Hollander AI, Bruchle NO, Zonneveld MN, Nurnberg G, Becker C, et al. Mutations of the CEP290 gene encoding a centrosomal protein cause Meckel-Gruber syndrome. Hum Mutat. 2008;29(1):45–52.

13. Leitch CC, Zaghloul NA, Davis EE, Stoetzel C, Diaz-Font A, Rix S, et al. Hypomorphic mutations in syndromic encephalocele genes are associated with Bardet-Biedl syndrome. Nat Genet. 2008;40(4):443–8.

14. Coppieters F, Lefever S, Leroy BP, and De Baere E. CEP290, a gene with many faces: mutation overview and presentation of CEP290base. Hum Mutat. 2010;31(10):1097–108.

15. Pazour GJ, and Witman GB. The vertebrate primary cilium is a sensory organelle. Curr Opin Cell Biol. 2003;15(1):105–10.

16. Satir P, and Christensen ST. Overview of structure and function of mammalian cilia. Annu Rev Physiol. 2007;69:377–400.

17. Singla V, and Reiter JF. The primary cilium as the cell’s antenna: signaling at a sensory organelle. Science. 2006;313(5787):629–33.

18. Yang TT, Su J, Wang WJ, Craige B, Witman GB, Tsou MF, et al. Superresolution Pattern Recognition Reveals the Architectural Map of the Ciliary Transition Zone. Sci Rep. 2015;5:14096.

19. Ringo DL. Flagellar motion and fine structure of the flagellar apparatus in Chlamydomonas. J Cell Biol. 1967;33(3):543–71.

20. Gilula NB, and Satir P. The ciliary necklace. A ciliary membrane specialization. J Cell Biol. 1972;53(2):494–509.

21. Sun S, Fisher RL, Bowser SS, Pentecost BT, and Sui H. Three-dimensional architecture of epithelial primary cilia. Proc Natl Acad Sci U S A. 2019;116(19):9370–9.

22. Robichaux M, Potter, VL, Zhang, Z, He, F, Schmid, MF, and Wensel, TG. Defining the Layers of a Sensory Cilium with STORM and Cryo-Electron Nanoscopies. 2019.

23. Pazour GJ, San Agustin JT, Follit JA, Rosenbaum JL, and Witman GB. Polycystin-2 localizes to kidney cilia and the ciliary level is elevated in orpk mice with polycystic kidney disease. Curr Biol. 2002;12(11):R378–80.

24. Handel M, Schulz S, Stanarius A, Schreff M, Erdtmann-Vourliotis M, Schmidt H, et al. Selective targeting of somatostatin receptor 3 to neuronal cilia. Neuroscience. 1999;89(3):909–26.

25. Gilliam JC, Chang JT, Sandoval IM, Zhang Y, Li T, Pittler SJ, et al. Threedimensional architecture of the rod sensory cilium and its disruption in retinal neurodegeneration. Cell. 2012;151(5):1029–41.

26. Salisbury JL. Centrin, centrosomes, and mitotic spindle poles. Curr Opin Cell Biol. 1995;7(1):39–45.

27. Wolfrum U. Centrin in the photoreceptor cells of mammalian retinae. Cell Motil Cytoskeleton. 1995;32(1):55–64.

28. Wolfrum U, and Salisbury JL. Expression of centrin isoforms in the mammalian retina. Exp Cell Res. 1998;242(1):10–7.

29. Laoukili J, Perret E, Middendorp S, Houcine O, Guennou C, Marano F, et al. Differential expression and cellular distribution of centrin isoforms during human ciliated cell differentiation in vitro. J Cell Sci. 2000;113 (Pt 8):1355–64.

30. Pulvermuller A, Giessl A, Heck M, Wottrich R, Schmitt A, Ernst OP, et al. Calcium-dependent assembly of centrin-G-protein complex in photoreceptor cells. Mol Cell Biol. 2002;22(7):2194–203.

31. Muresan V, and Besharse JC. Complex intermolecular interactions maintain a stable linkage between the photoreceptor connecting cilium axoneme and plasma membrane. Cell Motil Cytoskeleton. 1994;28(3):213–30.

32. Kim J, Krishnaswami SR, and Gleeson JG. CEP290 interacts with the centriolar satellite component PCM-1 and is required for Rab8 localization to the primary cilium. Hum Mol Genet. 2008;17(23):3796–805.

33. Tsang WY, Bossard C, Khanna H, Peranen J, Swaroop A, Malhotra V, et al. CP110 suppresses primary cilia formation through its interaction with CEP290, a protein deficient in human ciliary disease. Dev Cell. 2008;15(2):187–97.

34. Garcia-Gonzalo FR, Corbit KC, Sirerol-Piquer MS, Ramaswami G, Otto EA, Noriega TR, et al. A transition zone complex regulates mammalian ciliogenesis and ciliary membrane composition. Nat Genet. 2011;43(8):776–84.

35. Dharmat R, Eblimit A, Robichaux MA, Zhang Z, Nguyen TT, Jung SY, et al. SPATA7 maintains a novel photoreceptor-specific zone in the distal connecting cilium. J Cell Biol. 2018;217(8):2851–65.

36. Cox G, and Sheppard CJ. Practical limits of resolution in confocal and non-linear microscopy. Microsc Res Tech. 2004;63(1):18–22.

37. Gustafsson MG. Surpassing the lateral resolution limit by a factor of two using structured illumination microscopy. J Microsc. 2000;198(Pt 2):82–7.

38. Kim D, Deerinck TJ, Sigal YM, Babcock HP, Ellisman MH, and Zhuang X. Correlative stochastic optical reconstruction microscopy and electron microscopy. PLoS One. 2015;10(4):e0124581.

39. Graser S, Stierhof YD, Lavoie SB, Gassner OS, Lamla S, Le Clech M, et al. Cep164, a novel centriole appendage protein required for primary cilium formation. J Cell Biol. 2007;179(2):321–30.

40. Piperno G, and Fuller MT. Monoclonal antibodies specific for an acetylated form of alpha-tubulin recognize the antigen in cilia and flagella from a variety of organisms. J Cell Biol. 1985;101(6):2085–94.

41. Molday RS, and Molday LL. Identification and characterization of multiple forms of rhodopsin and minor proteins in frog and bovine rod outer segment disc membranes. Electrophoresis, lectin labeling, and proteolysis studies. J Biol Chem. 1979;254(11):4653–60.

42. Yang TT, Chong WM, Wang WJ, Mazo G, Tanos B, Chen Z, et al. Superresolution architecture of mammalian centriole distal appendages reveals distinct blade and matrix functional components. Nat Commun. 2018;9(1):2023.

43. Craige B, Tsao CC, Diener DR, Hou Y, Lechtreck KF, Rosenbaum JL, et al. CEP290 tethers flagellar transition zone microtubules to the membrane and regulates flagellar protein content. J Cell Biol. 2010;190(5):927–40.

44. Chang B. Mouse Models as Tools to Identify Genetic Pathways for Retinal Degeneration, as Exemplified by Leber’s Congenital Amaurosis. Methods Mol Biol. 2016;1438:417–30.

45. Cideciyan AV, Rachel RA, Aleman TS, Swider M, Schwartz SB, Sumaroka A, et al. Cone photoreceptors are the main targets for gene therapy of NPHP5 (IQCB1) or NPHP6 (CEP290) blindness: generation of an all-cone Nphp6 hypomorph mouse that mimics the human retinal ciliopathy. Hum Mol Genet. 2011;20(7):1411–23.

46. Chang B, Khanna H, Hawes N, Jimeno D, He S, Lillo C, et al. In-frame deletion in a novel centrosomal/ciliary protein CEP290/NPHP6 perturbs its interaction with RPGR and results in early-onset retinal degeneration in the rd16 mouse. Hum Mol Genet. 2006;15(11):1847–57.

47. Stone EM, Cideciyan AV, Aleman TS, Scheetz TE, Sumaroka A, Ehlinger MA, et al. Variations in NPHP5 in patients with nonsyndromic leber congenital amaurosis and Senior-Loken syndrome. Arch Ophthalmol. 2011;129(1):81–7.

48. Otto EA, Loeys B, Khanna H, Hellemans J, Sudbrak R, Fan S, et al. Nephrocystin-5, a ciliary IQ domain protein, is mutated in Senior-Loken syndrome and interacts with RPGR and calmodulin. Nat Genet. 2005;37(3):282–8.

49. Estrada-Cuzcano A, Koenekoop RK, Coppieters F, Kohl S, Lopez I, Collin RW, et al. IQCB1 mutations in patients with leber congenital amaurosis. Invest Ophthalmol Vis Sci. 2011;52(2):834–9.

50. Barbelanne M, Song J, Ahmadzai M, and Tsang WY. Pathogenic NPHP5 mutations impair protein interaction with Cep290, a prerequisite for ciliogenesis. Hum Mol Genet. 2013;22(12):2482–94.

51. Barbelanne M, Hossain D, Chan DP, Peranen J, and Tsang WY. Nephrocystin proteins NPHP5 and Cep290 regulate BBSome integrity, ciliary trafficking and cargo delivery. Hum Mol Genet. 2015;24(8):2185–200.

52. Basinger S, Bok D, and Hall M. Rhodopsin in the rod outer segment plasma membrane. J Cell Biol. 1976;69(1):29–42.

53. Heitzmann H. Rhodopsin is the predominant protein of rod outer segment membranes. Nat New Biol. 1972;235(56):114.

54. Lancaster MA, Gopal DJ, Kim J, Saleem SN, Silhavy JL, Louie CM, et al. Defective Wnt-dependent cerebellar midline fusion in a mouse model of Joubert syndrome. Nat Med. 2011;17(6):726–31.

55. Hynes AM, Giles RH, Srivastava S, Eley L, Whitehead J, Danilenko M, et al. Murine Joubert syndrome reveals Hedgehog signaling defects as a potential therapeutic target for nephronophthisis. Proc Natl Acad Sci U S A. 2014;111(27):9893–8.

56. Datta P, Hendrickson, B, Brendalen, S, Ruffcorn, A, and Seo, S. CEP290 myosin-tail homology domain is essential for protein confinement between inner and outer segments in photoreceptors. 2019.

57. Awata J, Takada S, Standley C, Lechtreck KF, Bellve KD, Pazour GJ, et al. NPHP4 controls ciliary trafficking of membrane proteins and large soluble proteins at the transition zone. J Cell Sci. 2014;127(Pt 21):4714–27.

58. Gogendeau D, Lemullois M, Le Borgne P, Castelli M, Aubusson-Fleury A, Arnaiz O, et al. MKS-NPHP module proteins control ciliary shedding at the transition zone. PLoS Biol. 2020;18(3):e3000640.

59. Megaw RD, Soares DC, and Wright AF. RPGR: Its role in photoreceptor physiology, human disease, and future therapies. Exp Eye Res. 2015;138:32–41.

60. Li T. Leber congenital amaurosis caused by mutations in RPGRIP1. Cold Spring Harbor perspectives in medicine. 2014;5(4).

61. Shu X, Fry AM, Tulloch B, Manson FD, Crabb JW, Khanna H, et al. RPGR ORF15 isoform co-localizes with RPGRIP1 at centrioles and basal bodies and interacts with nucleophosmin. Hum Mol Genet. 2005;14(9):1183–97.

62. Lau L, Lee YL, Sahl SJ, Stearns T, and Moerner WE. STED microscopy with optimized labeling density reveals 9-fold arrangement of a centriole protein. Biophys J. 2012;102(12):2926–35.

63. Tillberg PW, Chen F, Piatkevich KD, Zhao Y, Yu CC, English BP, et al. Protein-retention expansion microscopy of cells and tissues labeled using standard fluorescent proteins and antibodies. Nat Biotechnol. 2016;34(9):987–92.

64. Chang JB, Chen F, Yoon YG, Jung EE, Babcock H, Kang JS, et al. Iterative expansion microscopy. Nat Methods. 2017;14(6):593–9.

65. Macaluso FP, Perumal GS, Kolstrup J, and Satir P. CLEM Methods for Studying Primary Cilia. Methods Mol Biol. 2016;1454:193–202.

66. Norris RP, Baena V, and Terasaki M. Localization of phosphorylated connexin 43 using serial section immunogold electron microscopy. J Cell Sci. 2017;130(7):1333–40.

67. Lopes CA, Prosser SL, Romio L, Hirst RA, O’Callaghan C, Woolf AS, et al. Centriolar satellites are assembly points for proteins implicated in human ciliopathies, including oral-facial-digital syndrome 1. J Cell Sci. 2011;124(Pt 4):600–12.

68. Stowe TR, Wilkinson CJ, Iqbal A, and Stearns T. The centriolar satellite proteins Cep72 and Cep290 interact and are required for recruitment of BBS proteins to the cilium. Mol Biol Cell. 2012;23(17):3322–35.

69. Datta P, Hendrickson B, Brendalen S, Ruffcorn A, and Seo S. The myosin-tail homology domain of centrosomal protein 290 is essential for protein confinement between the inner and outer segments in photoreceptors. J Biol Chem. 2019;294(50):19119–36.

70. Drivas TG, Holzbaur EL, and Bennett J. Disruption of CEP290 microtubule/membrane-binding domains causes retinal degeneration. J Clin Invest. 2013;123(10):4525–39.

